# Vascular Endothelial-derived SPARCL1 Exacerbates Viral Pneumonia Through Pro-Inflammatory Macrophage Activation

**DOI:** 10.1101/2023.05.25.541966

**Authors:** Gan Zhao, Maria E. Gentile, Lulu Xue, Christopher V. Cosgriff, Aaron I. Weiner, Stephanie Adams-Tzivelekidis, Joanna Wong, Xinyuan Li, Sara Kass-Gergi, Nicolas P. Holcomb, Maria C. Basal, Kathleen M. Stewart, Joseph D. Planer, Edward Cantu, Jason D. Christie, Maria M. Crespo, Michael J. Mitchell, Nuala J. Meyer, Andrew E. Vaughan

## Abstract

Inflammation upon infectious lung injury is a double-edged sword: while tissue-infiltrating immune cells and cytokines are necessary to control infection, these same factors often aggravate injury. Full appreciation of both the sources and targets of inflammatory mediators is required to facilitate strategies to maintain antimicrobial effects while minimizing off-target epithelial and endothelial damage. Recognizing that the vasculature is centrally involved in tissue responses to injury and infection, we observed that pulmonary capillary endothelial cells (ECs) exhibit dramatic transcriptomic changes upon influenza injury punctuated by profound upregulation of *Sparcl1*. Endothelial deletion and overexpression of SPARCL1 implicated this secreted matricellular protein in driving key pathophysiologic symptoms of pneumonia, which we demonstrate result from its effects on macrophage polarization. SPARCL1 induces a shift to a pro-inflammatory “M1-like” phenotype (CD86^+^CD206^-^), thereby increasing associated cytokine levels. Mechanistically, SPARCL1 acts directly on macrophages *in vitro* to induce the pro-inflammatory phenotype via activation of TLR4, and TLR4 inhibition *in vivo* ameliorates inflammatory exacerbations caused by endothelial *Sparcl1* overexpression. Finally, we confirmed significant elevation of SPARCL1 in COVID-19 lung ECs in comparison with those from healthy donors. Survival analysis demonstrated that patients with fatal COVID-19 had higher levels of circulating SPARCL1 protein compared to those who recovered, indicating the potential of SPARCL1 as a biomarker for prognosis of pneumonia and suggesting that personalized medicine approaches might be harnessed to block SPARCL1 and improve outcomes in high-expressing patients.

## Introduction

Respiratory viral pathogens such as H1N1 and H5N1 influenza and SARS / SARS-CoV-2 can destroy alveolar epithelium both by direct infection and indirectly via cytokine release from infected cells, especially by type I and type III interferons (1, 2). In some patients this results in diffuse alveolar damage, impaired gas exchange, and ultimately acute respiratory distress syndrome (ARDS), which bears a mortality rate of >40% (3-5). Upon infection, activated immune cells release various cytokines, and although an appropriate inflammatory cytokine environment facilitates recruitment of immune cells for pathogen clearance and alveolar regeneration (6, 7), excessive accumulation of these cytokines can increase vascular permeability, induce additional cell death, and ultimately exacerbate lung injury (8, 9). This "cytokine storm", similar to Cytokine Release Syndrome seen after some CAR-T therapies, ultimately manifests as sepsis, a life-threatening complication that can present far out of proportion to the initial infection (10-12). Cytokine blocking strategies have been proposed to alleviate excessive inflammation, which may benefit virus-induced ARDS patients (13, 14), though clinical trials targeting various cytokines have often yielded underwhelming results. Given that morbidity and mortality from viral pneumonia are associated with excessive inflammation but strategies to control this remain limited, continued research is necessary to maintain the beneficial aspects of immune responses to viral agents while limiting unnecessary tissue damage.

Macrophages are the most abundant immune cells in the healthy lung and constitute the first line of defense of the respiratory system by recognizing and engulfing pathogens, releasing cytokines, and later, promoting tissue repair, making them a critical arm of the innate immune system (15). Macrophages exhibit remarkable phenotypic plasticity, adapting to microenvironmental cues and transforming into distinct phenotypes with specific functions. Activated macrophages are typically classified into two categories, M1 and M2, though these terms are somewhat controversial given widespread recognition of the fact that macrophage activation occurs along a spectrum rather than into discrete subtypes. As an extension of the “Th1/Th2” paradigm of immune responses, M1 macrophages are defined as macrophages that produce pro-inflammatory cytokines and mediate resistance to intracellular pathogens, but these also lead to tissue destruction. M2 macrophages are in turn involved in anti-inflammatory responses and tissue repair / remodeling (16). Altering the activation state of macrophages to better control the inflammatory environment represents a promising strategy for the treatment of various diseases.

The "state" of macrophage activation is regulated by a complex set of signals. Accumulated evidence indicates that endothelial cells (ECs) lining the pulmonary vasculature regulate lung function not only through oxygen and nutrient delivery, but also participate in alveolar regeneration, immune responses, and fibrotic remodeling through production and release of paracrine signals, also known as angiocrine factors (17-21). Prior work has identified EC-derived signals that alter macrophage activation state to protect the lungs from injury (22, 23). Secreted protein acidic and rich in cysteine-like protein 1 (SPARCL1), a matricellular protein reported to inhibit angiogenesis in colorectal carcinoma but also contribute to nonalcoholic steatohepatitis progression in mice (24, 25), exhibits greatly increased expression after influenza injury. However, whether SPARCL1 contributes to viral pneumonia progression is unknown. Here, we demonstrate lung capillary ECs adopt a distinct transcriptomic and phenotypic state upon viral injury to generate high levels of SPARCL1, which in turn acts through TLR4 to promote a pro-inflammatory M1-like state in macrophages, exacerbating inflammation and increasing the severity of viral pneumonia. Further, we show that antagonism of TLR4 can specifically rescue morbidity in SPARCL1 overexpressing mice, suggesting a potential therapeutic intervention for pneumonia patients with high levels of circulating SPARCL1.

## Results

### Dynamic endothelial transcriptomics reveal increased Sparcl1 expression after influenza injury

Pulmonary gas exchange restoration after viral pneumonia requires vascular repair to restore the heterogeneous assemblage of lung endothelial cells (ECs) (9, 26, 27). To explore the dynamics of endothelial subpopulations during regeneration after viral lung injury, mouse lung ECs were isolated (CD45^-^EpCAM^-^CD31^+^) by FACS on day 0 (D0, uninjured), day 20 (D20) and day 30 (D30) post influenza infection, and subsets/clusters were then identified by single-cell transcriptomic profiling (**Fig.1A**). Based on the well characterized EC subset signature genes (28, 29), we identified 6 EC clusters (**Suppl Fig.1A-B**), including lymphatic ECs (*Prox1*), venous ECs (*Bst1*), arterial ECs (*Gja5*), proliferating ECs (*Mki67*), and 2 capillary EC clusters, aerocytes (aCap, *Car4,*) and general capillary ECs (gCap, *Gpihbp1*). Further analysis of these EC subtypes revealed 2 lymphatic endothelial subsets, lymphatic ECs_01 (Ccl21a^hi^ lym_ECs, signature genes: *Ccl21a*, *Mmrna*, and *Nts*) and lymphatic ECs_02 (Prox1^hi^ lym_ECs, signature genes: *Prox1*, *Sned1*, and *Stab1*), and 3 gCap subsets, gCap ECs_01 (Dev.ECs, signature genes: *Hpgd*, *Tmem100*, and *Atf3*), gCap ECs_02 (Immu.EC, signature genes: *Cd74*, *Sparcl1* and *Cxcl12*) and gCap ECs_03 (*Gm26917*, *Nckap5*, and *Syne1*) (**Suppl Fig.1C**). GO (gene ontology) pathway enrichment analysis revealed gCap ECs_01 and gCap ECs_02 genes were enriched in vascular developmental and immune regulatory pathways respectively (**Suppl Fig.1D**), confirming the identification of gCap ECs, Dev.ECs (devEC) and Immu.ECs (immuneEC) recently reported by Zhang et al.(27).

**Fig.1.**
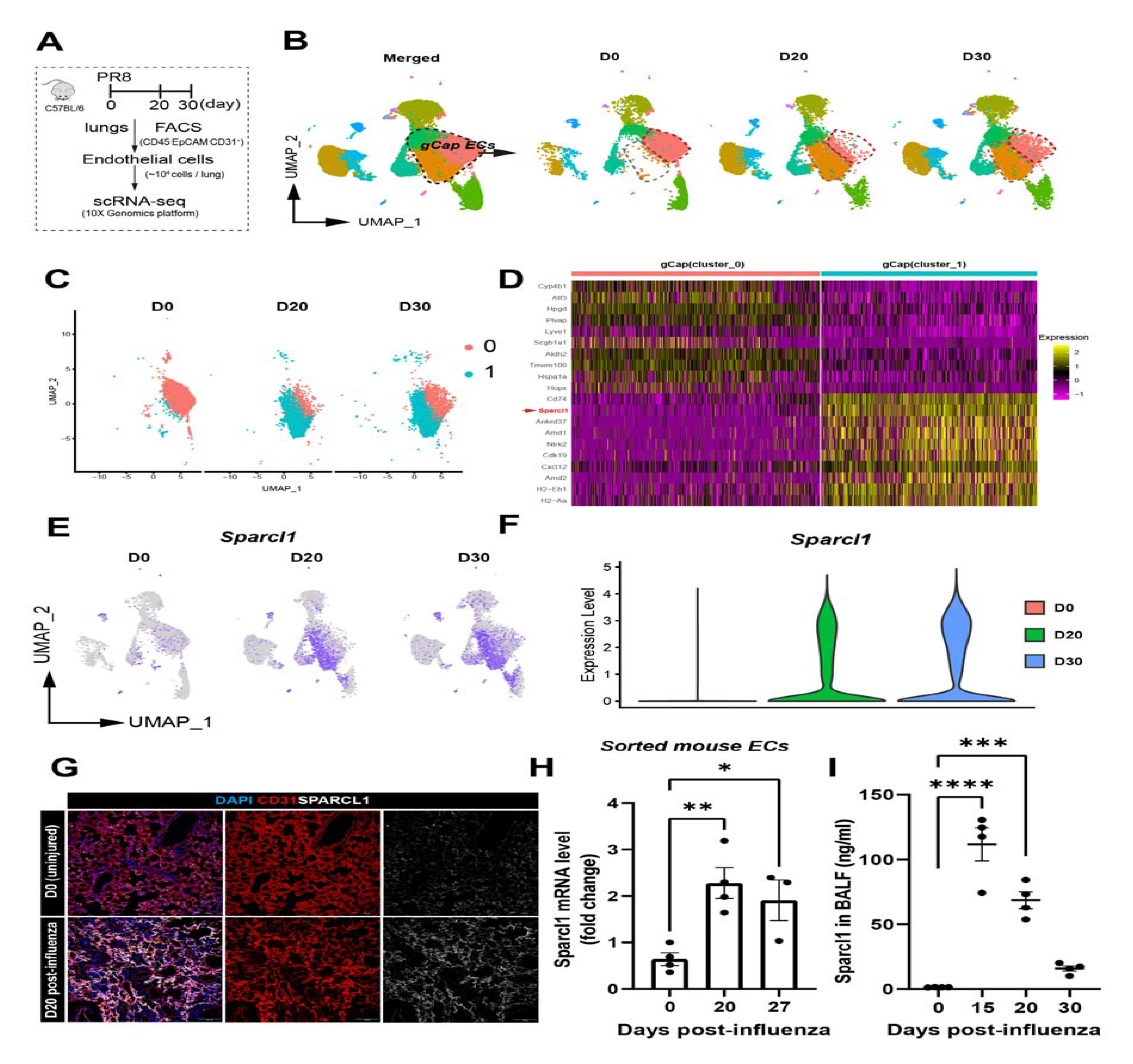
Single-cell transcriptomes reveal transcriptional dynamics in gCap ECs and increased SPARCL1 expression after viral injury. **A.** Schematic of mouse lung endothelial single-cell sequencing preparation. **B.** ScRNA-seq analysis for mouse lung ECs sorted from uninjured (D0) and on 20 and 30 days after influenza infection (marked as D20 and D30, respectively). Uniform manifold approximation and projection (UMAP) plots showing the dynamics in gCap ECs. **C.** The 2 gCap EC clusters of interest, cluster_0 (Dev.ECs) and cluster_1 (Immu.ECs) were subsetted from (**B**). **D.** Heatmap showing the top 20 differentially expressed genes of cluster_0 (Dev.ECs) and cluster_1(Immu.ECs). **E.** UMAP analysis reveals that *Sparcl1* is predominantly expressed in gCap ECs, especially in Immu.ECs. **F.** Violin plots showing *Sparcl1* expression level in mouse lung ECs sorted from D0, D20 and D30, respectively. **G.** Representative immunostaining of SPARCL1 in endothelial cells (CD31) in both uninjured (D0) and D20 after influenza infection lung tissues. Scale bar, 100 μm. **H.** qPCR analysis of *Sparcl1* in isolated lung ECs (CD45^−^CD31^+^) sorted on days 0 (uninjured), 20 and 27 after influenza infection, n = 3-4 mice per group. **I.** The concentration of SPARCL1 in bronchoalveolar lavage fluid (BALF) was measured by ELISA at 0 (uninjured), 15, 20, and 30 days after influenza infection, n = 4 mice per group. Data in (**H**) and (**I**) are presented as means ± SEM, calculated using one-way analysis of variance (ANOVA), followed by Dunnett’s multiple comparison test. *P < 0.05, **P < 0.01, ***P < 0.001 and ****P < 0.0001.

Interestingly, we found that the two subgroups of gCap ECs, Dev.ECs and Immu.ECs, exhibited dynamic changes during injury, with Dev.ECs significantly reduced during infection (D20) and gradually returning to baseline levels after recovery from pneumonia (D30). Immu.EC, on the other hand, showed the opposite trend (**Fig.1B**). Largely absent in uninjured lungs, Immu.ECs mainly appeared after injury, indicating a potential role in vascular responses to injury. Therefore, we focused on transcriptomic changes in these two gCap EC subtypes (**Fig.1C**). SPARCL1, a matricellular protein reported to inhibit angiogenesis in colorectal carcinoma(25), was broadly expressed in Immu.ECs and significantly increased in ECs after injury (**Fig.1D-F**). Immunostaining showed that SPARCL1 was mainly expressed in capillary ECs (especially gCap ECs), as well as mesenchymal/stromal cells (**Suppl Fig.2A-C**), and was significantly increased on day 20 after influenza injury (**Fig.1G**). Subsequently, quantitative polymerase chain reaction (qPCR) analysis for *Sparcl1* mRNA in isolated ECs on day 0, 20 and 27 post influenza and enzyme-linked immunosorbent assay (ELISA) analysis for SPARCL1 protein in bronchoalveolar lavage fluid (BALF) collected on day 0, 15, 20 and 30 post influenza confirmed the increased *Sparcl1*/SPARCL1 expression after injury (**Fig.1H and I**). Taken together, these results demonstrate a subpopulation of gCap EC (Immu.ECs) expressing very high levels of *Sparcl1* appears during injury.

### Endothelial ablation of Sparcl1 mitigates influenza-induced lung injury

Given that the injury-induced population of Immu.ECs were characterized by high expression of *Sparcl1*, we proceed to further probe the function of this gene. To explore the role of endothelial SPARCL1 in the pathogenesis of viral pneumonia, we crossed VECad^CreERT2^ mice with novel Sparcl1^flox^ mice. We used a CRISPR-Cas9 strategy in mouse embryonic stem cells (**Fig.2A and B**) to generate homozygous mutant mice and, upon crossing, ultimately proceeded with selective ablation of Sparcl1 in ECs of adult mice (referred to as EC^Sparcl1-KO^) via tamoxifen administration (**Fig.2C**). Sparcl1^flox/flox^ mice lacking Cre (referred to as WT) were used as the control group. Intriguingly, EC^Sparcl1-KO^ mice demonstrated less severe pneumonia symptoms, evidenced by less initial body weight loss and faster recovery to baseline levels (**Fig.2D**) as well as improved capillary oxygen saturation compared to WT mice (**Fig.2E**). Additionally, EC^Sparcl1-KO^ mice trended toward higher probability of survival (∼75%) compared to WT (less than 50%) by day 27 post infection when challenged with a high dose of influenza virus (**Fig.2F**). We next analyzed local inflammatory cytokines associated with viral pneumonia, including respiratory distress induced by both influenza and SARS-Cov2 (11, 30, 31). Cytokine levels in the BALF were normal during homeostasis, but EC^Sparcl1-KO^ mice exhibited lower levels of the inflammatory cytokines TNF-α and IL-6 on day 12 after influenza infection compared with WT mice (**Fig.2G and H**). Moreover, we also observed decreased total protein and cells in the bronchoalveolar lavage fluid (BALF) of EC^Sparcl1-KO^ mice on day 12 post infection (**Fig.2I and J**). Next, we assessed changes in the major immune cell populations in the lung and found that endothelial deletion of Sparcl1 significantly affected the number of recruited macrophages but not other immune cells including T, B, natural killer (NK) and innate lymphoid cells (ILCs) (**Fig.2K and Suppl Fig.3A-B**). These results indicate that mice lacking EC *Sparcl1* exhibit dampened local inflammatory responses, further corroborated histologically, as these lungs bore less inflammatory immune cell infiltration and more orderly alveolar structure and thickness in the area of injury on day 25 post-infection (**Fig.2L**). These experiments indicate that endothelial deficiency in Sparcl1 protects against severe influenza pneumonia, at least partially by attenuating local inflammation.

**Fig.2.**
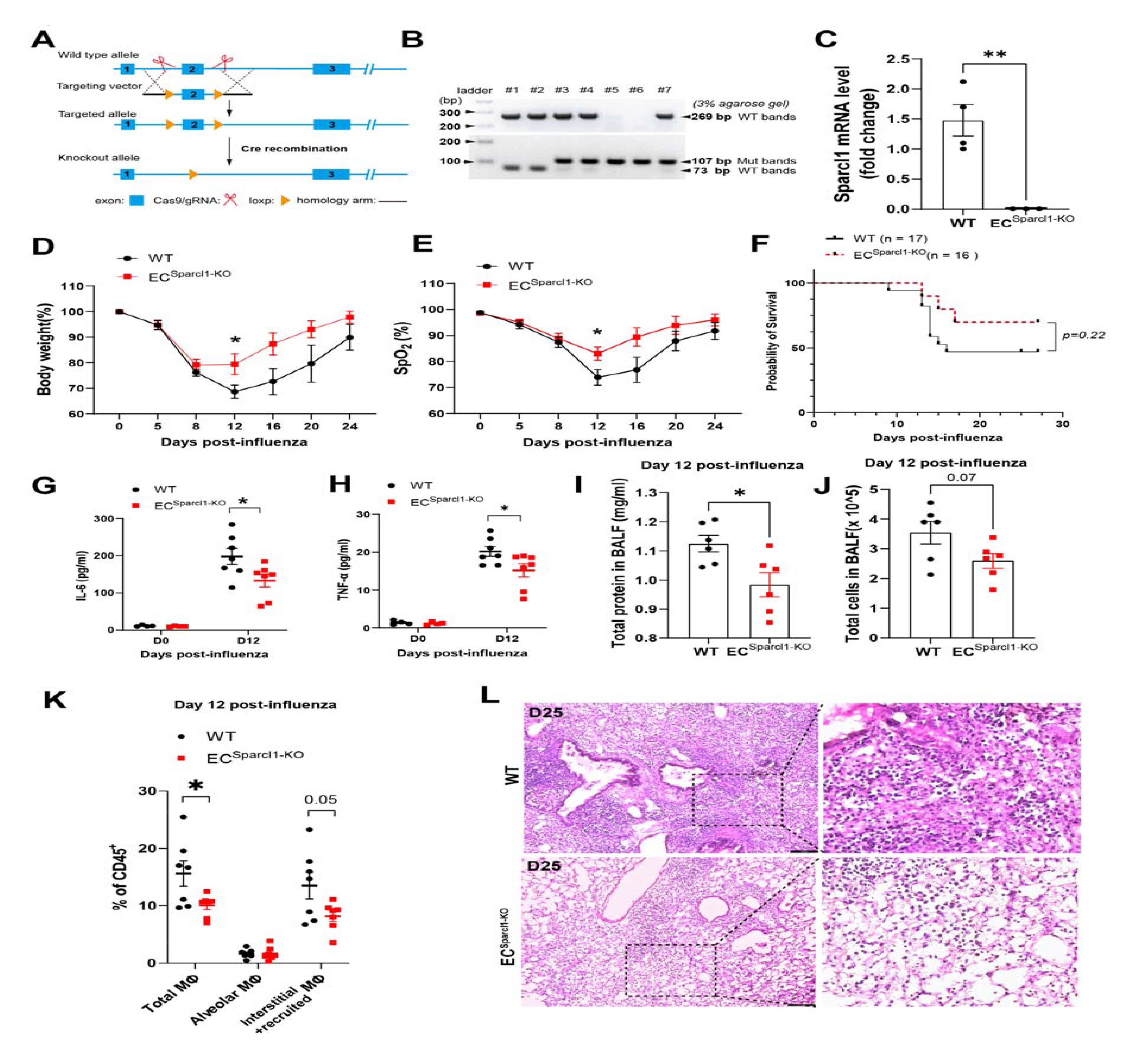
Endothelial loss of Sparcl1 attenuates influenza-induced pneumonia. **A.** Schematic of the strategy used to generate Sparcl1 loxP/+ mice using CRISPR/Cas9 technology. **B.** DNA gel image of Sparcl1^flox^ mouse genotyping. Lanes 1 and 2 represent wild-type mice with PCR product at 269 bp. Lanes 3, 4 and 7 represent heterozygous mice with product at 269 bp (wt) and product at 113 bp (Primers designed upstream and downstream of the loxp site). A light band at 79 bp from amplification of the WT locus can be observed as well in heterozygous mice but additional primers genotyping for WT allele (upper) is used to confirm. Lanes 5 and 6 represent homozygous mice, where LoxP sites were inserted at both alleles (single 119-bp PCR product). **C.** qPCR analysis of *Sparcl1* in isolated lung ECs (CD45^−^CD31^+^) sorted from EC^Sparcl1-KO^ (VECad^CreERT2^; Sparcl1^flox/flox^) or WT (Sparcl1^flox/flox^) mice 2 weeks after 5 doses of tamoxifen injection. **D-E**. (**D**)Time course of changes in body weight and (**E**) capillary oxygen saturation in WT and EC^Sparcl1-KO^ mice after influenza infection, n = 5 to 7 mice per group. **F**. Kaplan-Meier survival curves after influenza infection, log-rank test. **G-H**. The levels of pro-inflammatory cytokines IL-6 (**G**) and TNF-α (**H**) in bronchoalveolar lavage fluid (BALF) were measured by ELISA in WT and EC^Sparcl1-KO^ mice at day 0 (uninjured) and 12 days after influenza infection, n = 4 to 7 mice per group. **I-J.** Total protein (**I**) and cells (**J**) were quantified in BALF on day 12 post influenza infection, n = 6 to 7 per group. **K.** Quantification of the proportion of total macrophages (CD64^+^F4/80^+^), alveolar macrophages (CD45^+^Ly6G^-^CD64^+^F4/80^+^SiglecF^+^) and interstitial and recruited macrophages (CD45^+^Ly6G^-^CD64^+^F4/80^+^SiglecF^-^) in CD45^+^ live cells at day 12 after influenza infection in WT and EC^Sparcl1-KO^ mice, n = 6 to 7 mice per group. **L.** Representative histological changes in the lungs of influenza-challenged WT and EC^Sparcl1-KO^ mice at day 25 post-infection. Scale bars, 100 μm. Data in (**C**) to (**E**), (**G**) and (**J**) are presented as means ± SEM, calculated using unpaired two-tailed t test. Data in (**F**) were calculated using log-rank test. *P < 0.05, **P < 0.01.

### Sparcl1 overexpression worsens influenza-induced pneumonia

To further confirm that EC-derived SPARCL1 negatively contributes to pneumonia severity, we again targeted *Sparcl1* cDNA to the ROSA26 locus, preceded by a loxP flanked "stop" sequence, in mouse ES cells. Upon generation of these mice, we then crossed these animals with the VECad^CreERT2^ strain to develop conditional endothelial *Sparcl1* knock-in/overexpression mice (**Fig.3A and B**), VECad^CreERT2^; Sparcl1^+/WT^ or VECad^CreERT2^; Sparcl1^+/+^ mice (referred to as EC^Sparcl1-OE^). VECad^CreERT2^; Sparcl1^WT/WT^ mice (referred to as WT) were used as the control group. Upon tamoxifen administration, SPARCL1 overexpression in the lungs of EC^Sparcl1-OE^ mice was confirmed by western blotting (**Fig.3C**). As expected, upon influenza infection, EC^Sparcl1-OE^ mice displayed exaggerated pneumonia outcomes, as demonstrated by greater weight loss, prolonged recovery period (related to baseline level of body weight at D0) (**Fig.3D**), worse lung respiratory function during pneumonia (impaired oxygen saturation) (**Fig.3E**), and increased total protein and cells in the BALF (**Fig.3F and G**). To investigate whether overexpression of SPARCL1 exacerbates the local inflammatory response during pneumonia, we examined typical pro-inflammatory cytokines TNF-α, IL-1β, and IL-6 in BALF. As with *Sparcl1* deletion, no significant changes in these cytokines occurred during homeostasis, though they began to trend higher on day 10 post-infection in EC^Sparcl1-OE^ mice compared with WT mice. However, cytokine levels were significantly elevated in comparison to WT mice by day 20, likely contributing to the extended physiologic recovery time observed in EC^Sparcl1-OE,^ reflecting long-term inflammation (**Fig.3H**). Similarly, overexpression of SPARCL1 significantly increased the number of pulmonary macrophages, especially recruited/interstitial macrophages, but without obvious effects on other immune cells, including T, B, NK and ILC cells (**Fig.3I and Suppl Fig.3C**). This further supports the previous conclusion that EC-derived SPARCL1 aggravates local lung inflammation in pneumonia, thus worsening lung injury.

**Fig.3.**
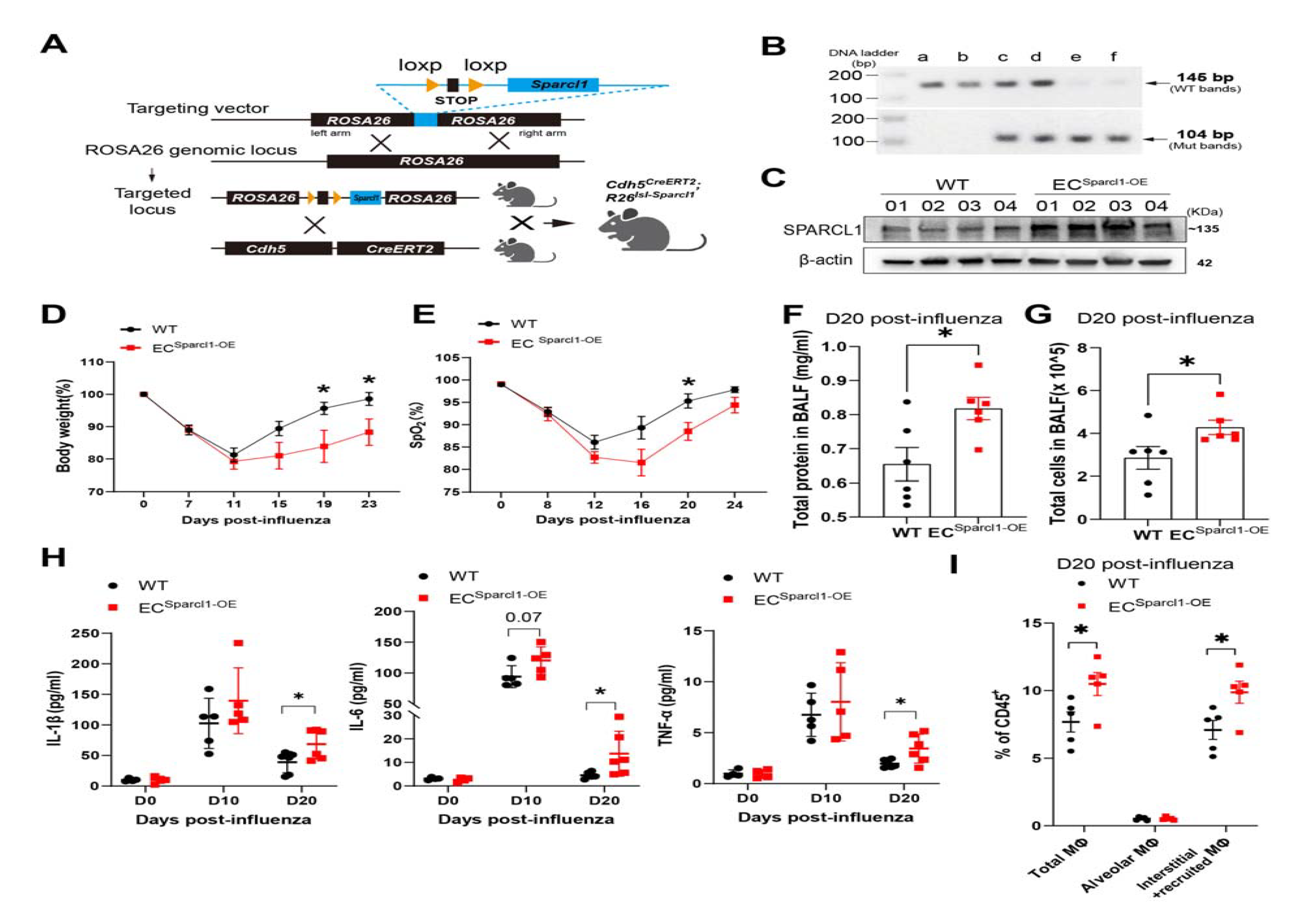
Endothelial overexpression of Sparcl1 exacerbates influenza-induced pneumonia. **A.** Schematic of the strategy used to generate Sparcl1 knock-in mice targeting ROSA26 locus. **B.** DNA gel image for genotyping of endothelial Sparcl1 knock-in mice. Lanes *a* and *b* represent wild-type mice with PCR product at 145 bp. Lanes *c* and *d* represent heterozygous mice with product at 145 bp (wt) and product at 104 bp (mut). Lanes *e* and *f* represent homozygous mice with PCR product at 104 bp. **C.** Western blot of SPARCL1 in whole lung tissue from EC^Sparcl1-OE^ (VECad^CreERT2^; Sparcl1^+/wt^ or VECad^CreERT2^; Sparcl1^+/+^) or WT (VECad^CreERT2^) mice 2 weeks after 5 doses of tamoxifen. **D-E**. (**D**) Time course of changes in body weight and (**E**) capillary oxygen saturation in WT and EC^Sparcl1-OE^ mice after influenza infection, n = 5 to 7 mice per group. **F-G.** Total protein (**F**) and cells (**G**) were quantified in BALF on day 20 post influenza infection, n = 6 mice per group. **H**. The concentration of pro-inflammatory cytokines, IL-1β, IL-6 and TNF-α in BALF were measured by ELISA in WT and EC^Sparcl1-OE^ mice at 0 (uninjured, n = 3/group), 10(n = 5/group) and 20 (n = 5/group) days post-influenza infection. **I.** Quantification of the proportion of total macrophages (CD64^+^F4/80^+^), alveolar macrophages (CD45^+^Ly6G^-^CD64^+^F4/80^+^SiglecF^+^) and interstitial and recruited macrophages (CD45^+^Ly6G^-^ CD64^+^F4/80^+^SiglecF^-^) in CD45^+^ live cells at day 20 after influenza infection in WT and EC^Sparcl1-OE^ mice, n = 5 mice per group. Data in (**D**) to (**G**) and (**I**)are presented as means ± SEM, Data in (**H)** is presented as means ± SD, calculated using unpaired two-tailed t test. *P < 0.05.

### SPARCL1 promotes proinflammatory changes in macrophage phenotypes *in vivo*

To further address the mechanisms underlying exacerbation of pneumonia outcomes attributed to endothelial Sparcl1, we assessed whether ablation or overexpression of Sparcl1 affected EC angiogenic proliferation, as predicted by reports that SPARCL1 can act as an angiostatic factor (25). We administered the nucleoside analog 5-ethynyl-2-deoxyuridine (EdU) (50 mg/kg, intraperitoneally) and intracellular EdU flow analysis was used to quantify proliferative ECs (**Suppl Fig.4A**). Surprisingly, we observed no significant changes in EC proliferation upon either endothelial ablation (**Suppl Fig.4A-C**) or overexpression (**Suppl Fig.4D-F**) of Sparcl1. Given these results and the fact that SPARCL1 is a secreted matricellular protein, we reasoned that SPARCL1 may instead be acting as a paracrine signaling molecule, influencing the phenotype of other cell types involved in lung injury and repair.

Since SPARCL1 triggered more severe inflammation upon injury and macrophage (MΦ) infiltration, (**Fig. 2F-L and 3F-I**), we speculated that SPARCL1 promotes release of pro-inflammatory cytokines by affecting macrophage polarization. Macrophages participate in both injury / inflammation and tissue repair by releasing cytokines and chemokines and can be subdivided into classical activated (M1) and alternatively activated (M2) macrophages which exhibit pro-and anti-inflammatory phenotypes, respectively. M1 macrophages are reported as the main sources of several proinflammatory cytokines, including TNF-α, IL-1β and IL-6, acting to amplify the inflammatory response (32, 33). Thus, we examined whether SPARCL1 contributes to pro-inflammatory responses by affecting macrophage M1/M2 polarization. Lung macrophages (CD45^+^Ly6G^-^CD64^+^F4/80^+^) were sub-gated into SiglecF^+^ alveolar macrophages (AMs) and SiglecF^-^ interstitial and/or recruited macrophages (I/RMs). Macrophages were then sub-phenotyped using the well-described CD86 and CD206 antigens(34) to distinguish M1_like (CD86^+^CD206^-^) and M2_like (CD86^-^CD206^+^) macrophage populations *in vivo* (**Suppl Fig.5A**). We observed significantly increased M1_like and decreased M2_like macrophage populations in EC^Sparcl1-OE^ mice compared with WT mice on day 20 post influenza infection, both in total MΦ (CD64^+^F4/80^+^) as well as specifically within AMs and I/RMs (**Fig.4A-C, Suppl Fig.5A**). However, EC overexpression of Sparcl1 did not significantly alter the M1/M2 macrophage proportions under homeostasis (**Suppl Fig.5B**), which we postulate is due to SPARCL1 protein failing to infiltrate into the alveolar space when the blood vessels are intact without injury. Immunostaining for the M2_like macrophage marker RELMα showed a decreased proportion of M2_like macrophages (RELMα^+^F4/80^+^ / F4/80^+^) in EC^Sparcl1-OE^ mice *in situ* on day 20 post-influenza infection (**Fig.4D**), further confirming our flow cytometry data.

**Fig.4.**
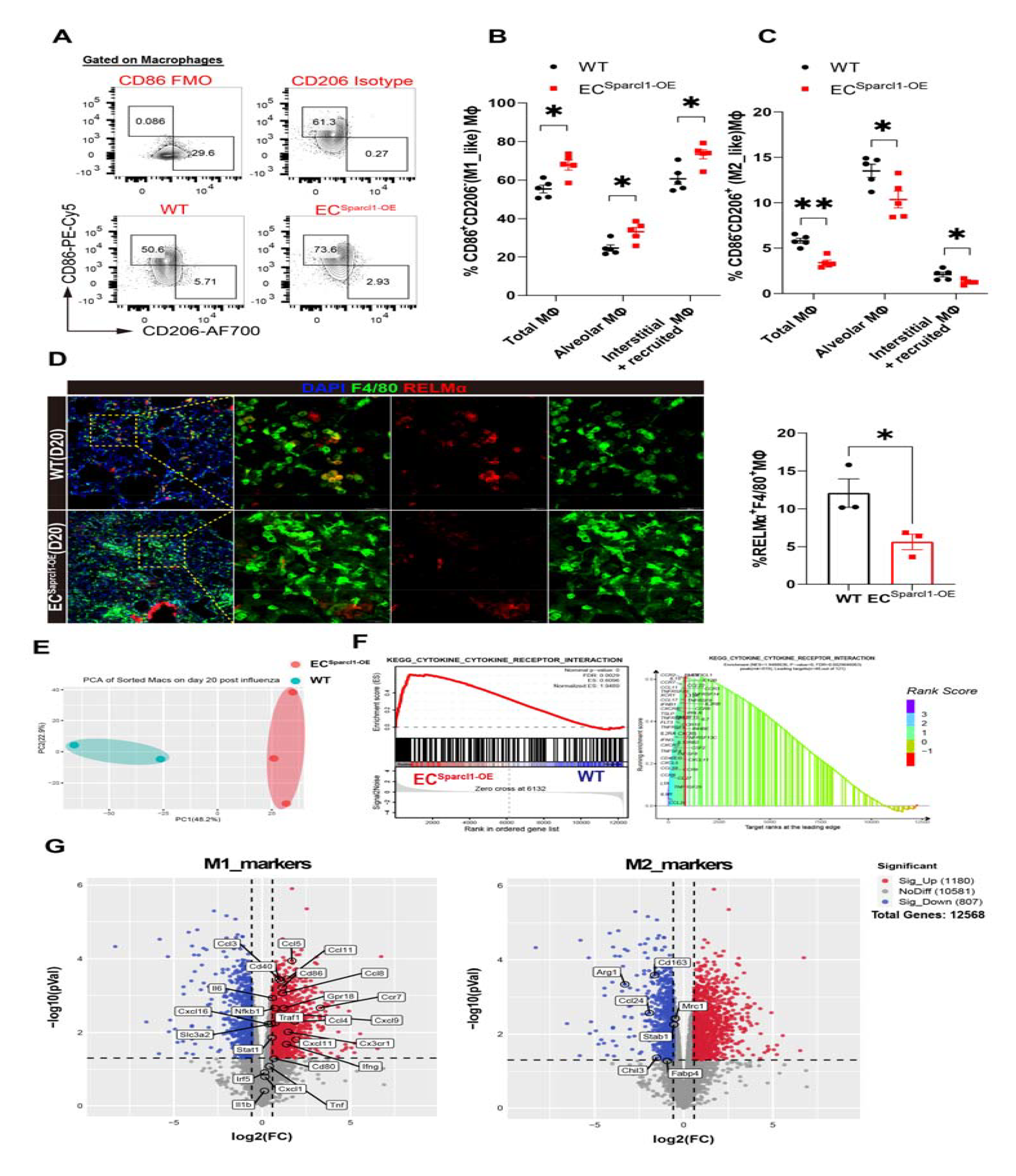
Endothelial overexpression of Sparcl1 promotes M1-like polarization of lung macrophages. **A.** Representative gating scheme for identification of pulmonary M1-like (CD86^+^CD206^-^) and M2-like (CD86^-^CD206^+^) macrophages (CD45^+^Ly6G^-^CD64^+^F4/80^+^) at day 20 after influenza infection in WT and EC^Sparcl1-OE^ mice. **B-C**. Quantification of the proportion of (**B**) M1-like (CD86^+^CD206^-^) and (**C**) M2-like (CD86^-^ CD206^+^) macrophages in total lung macrophages (CD45^+^Ly6G^-^CD64^+^F4/80^+^), alveolar macrophages (CD45^+^Ly6G^-^CD64^+^F4/80^+^SiglecF^+^) and interstitial and recruited macrophages (CD45^+^Ly6G^-^CD64^+^F4/80^+^SiglecF^-^) at day 20 after influenza infection in WT and EC^Sparcl1-OE^ mice, n = 5 mice per group. **D.** Left: representative immunostaining of lung M2_like macrophage (RELMα^+^F4/80^+^) in WT and EC^Sparcl1-OE^ mice on day 20 post influenza infection, scale bars, 100 μm (merge). 25 μm (separate chanel). Right: quantification of the proportion of M2_like macrophages (RELMα^+^F4/80^+^ / F4/80^+^) in left, n = 3 mice per group. **E.** Principal components analysis (PCA) indicates transcriptomic changes in lung macrophages between WT and EC^Sparcl1-OE^ mice on day 20 post influenza infection. **F.** Gene Set Enrichment Analysis (GSEA) of RNA-seq profiles of purified lung macrophages isolated from WT and EC^Sparcl1-OE^ mice on day 20 post influenza infection. Enrichment score and p value are displayed. **G.** Volcano plot indicating the M1-associated genes upregulated in lung macrophage from EC^Sparcl1-OE^ mice and M2-associated genes down-regulated on day 20 post influenza infection. Each point represents one mouse. Data in (**B**) and (**D**) are presented as means ± SEM, calculated using unpaired two-tailed t test. *P < 0.05, **P < 0.01.

RNA sequencing (RNA-seq) of isolated lung macrophages (CD45^+^Ly6G^-^CD64^+^F4/80^+^) on day 20 post-influenza infection was used to further examine the consequences of EC overexpression of Sparcl1. Principal components analysis (PCA) analysis was performed, with EC^Sparcl1-OE^ samples forming a tight cluster distinct from WT samples, confirming that EC Sparcl1 overexpression results in significant transcriptional variation effects on macrophages (**Fig.4E**). GSEA (gene set enrichment analysis) demonstrated differentially expressed genes were significantly enriched in the cytokine-cytokine receptor signaling pathway, and specifically, EC^Sparcl1-OE^ positively regulated the expression of inflammatory pathway genes (cytokines, receptors, etc.) (**Fig.4F**). Differential gene analysis showed that typical M1_like signature genes, including *Cd86*, *Cd80*, and *Ifng*, *Ccl3*, *Ccl4*, *Ccl5*, etc were significantly up-regulated in EC^Sparcl1-OE^ mouse macrophages, while M2_like signature genes, including *Arg1*, *Cd163*, *Mrc1*, and *Chil3* were significantly down-regulated (**Fig.4G**), suggesting that the macrophage polarization switch is a direct response to endothelial overexpression of Sparcl1. We also asked whether endothelial deletion of Sparcl1 might inversely affect M1/M2-like macrophage polarization during injury in comparison to Sparcl1 overexpression. Macrophages and their M1/M2 subsets were again gated as mentioned above on day 12 post influenza infection (**Suppl Fig.6A**). In agreement with results using EC^Sparcl1-OE^ mice, EC loss of Sparcl1 resulted in fewer M1_like but more M2_like macrophages during pneumonia (**Suppl Fig.6B-D**), with no obvious changes under homeostasis (**Suppl Fig.6E and F**). Taken together, these data support a model wherein EC-derived SPARCL1 induces an inflammatory response by triggering the polarization of M1-like macrophages and/or inhibiting the transformation of M2 macrophages, thereby promoting the persistence of inflammation that contributes to the exacerbation of pneumonia.

### The SPARCL1 induced pro-inflammatory macrophage phenotype requires TLR4 signaling ***in vitro***

To better understand the molecular basis of SPARCL1 induced M1 macrophage polarization, bone marrow (BM) cells were isolated and differentiated into mature macrophages, BM derived macrophages (BMDMs), which were validated by flow cytometry analysis (CD11b^+^F4/80^+^) (**Suppl Fig.7A-B**) and then treated with 5, 10, 20 μg/ml recombinant mouse SPARCL1 protein. NF-κB acts as the critical M1 polarization regulator via transcriptional activation of pro-inflammatory cytokine gene expression(16), and SPARCL1 significantly induced phosphorylation of NF-κB p65 (**Fig.5A**) and the release of pro-inflammatory cytokines TNF-α, IL-1β and IL-6 (**Fig.5B**). Morphologically, SPARCL1-treated BMDMs appeared similar to BMDMs treated with lipopolysaccharide (LPS, a known activator of the M1 phenotype), further indicating that SPARCL1 induces BMDMs towards an M1 pro-inflammatory phenotype (**Fig.5C**). Next, we explored whether SPARCL1 enhances an M1_like phenotype or hinders the M2_like phenotype. BMDMs were polarized into M1/M2 macrophages by treatment with LPS (50 ng/ml) or IL-4 (20 ng/ml) for 24 hours, and subsequently incubated with recombinant SPARCL1 protein (10 μg/ml) for 24 hours (**Fig.5D**). Our data showed that M2_like macrophages (CD206^+^CD11b^+^F4/80^+^) retain significant CD206 expression even upon withdrawal of IL-4 treatment for 24 hours (M2+PBS, ∼80%), but this was significantly reduced by SPARCL1 treatment (**Fig.5E and F**). Further analysis of the supernatant from these experiments demonstrated that IL-1β and IL-6 levels were slightly increased after SPARCL1 treatment compared with M1 control group (**Suppl Fig.7C**), however, SPARCL1 significantly increased the levels of TNFα, IL-1β and IL-6 in comparison with M2 control (**Suppl Fig.7C**). Moreover, the M2 macrophage marker genes *Mrc1* and *Chil3* were detected by qPCR and were highly expressed in the M2 control group as expected but suppressed upon subsequent SPARCL1 treatment (**Fig.5G**). M2 macrophages treated with SPARCL1 were morphologically distinct from the M2 control, again exhibiting a pro-inflammatory M1 macrophage appearance (**Suppl Fig.7D**). These data indicate that SPARCL1 suppresses the M2_like phenotype and promotes a transition toward the M1 proinflammatory phenotype.

**Fig.5.**
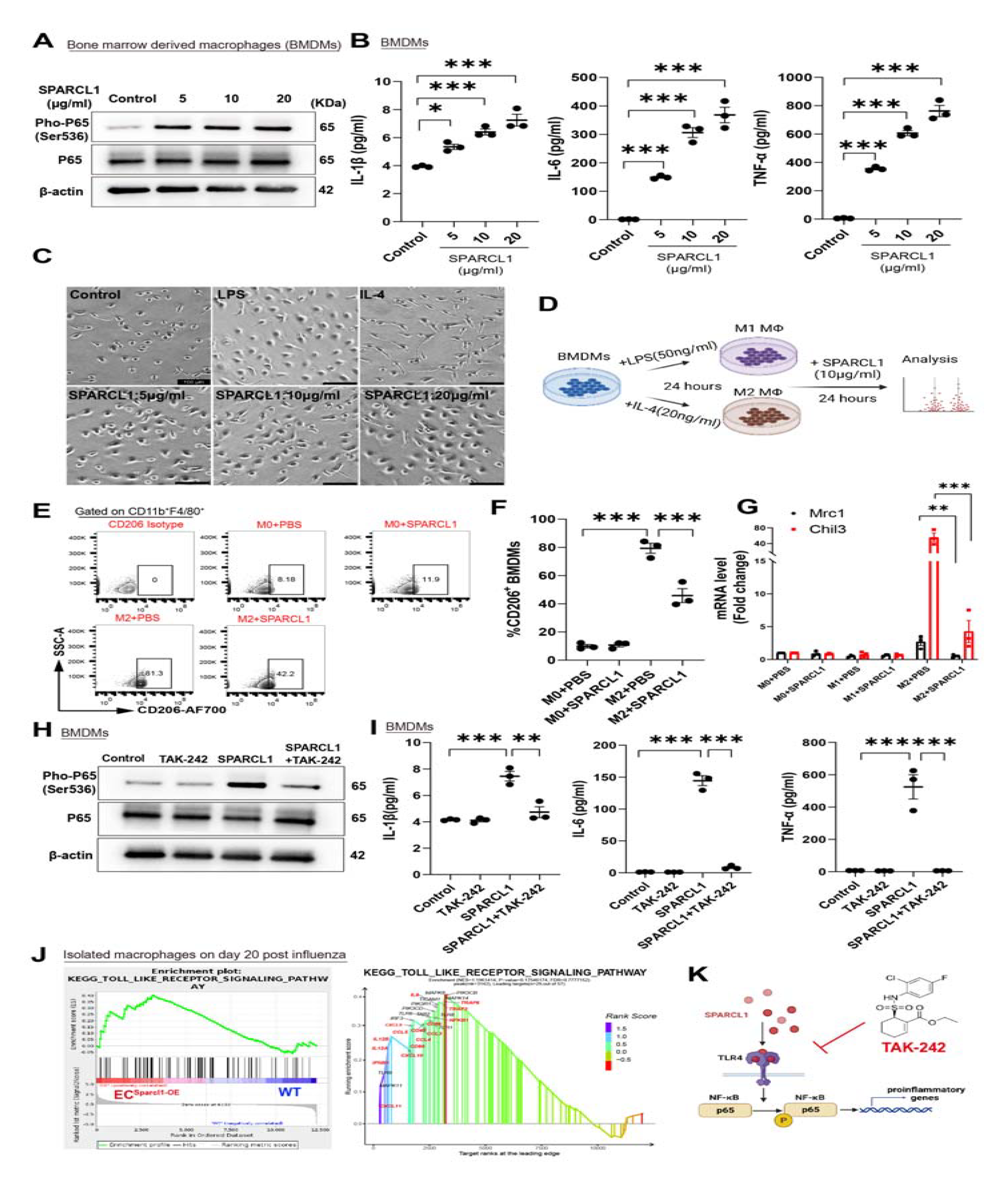
Sparcl1 induction of M1-like macrophages depends on TLR4 *in vitro*. **A.** Western blotting analysis of indicated proteins in bone marrow derived macrophages (BMDMs) treated with different doses of recombinant SPARCL1 protein (0-20 μg/ml) for 1 hour. **B.** BMDMs were treated with different doses of recombinant SPARCL1 protein (0-20 μg/ml) for 24 hours, and cell supernatant was collected. Th1 cytokines IL-1β, IL-6 and TNF-α levels in supernatant were measured by ELISA, n = 3 per group. **C.** BMDMs were treated with lipopolysaccharides (LPS, 50 ng/ml), IL-4 (20 ng/ml), SPARCL1 (5, 10 and 20 μg/ml), or vehicle control for 24 hours. Cell morphology was observed and imaged by light microscopy. Scale bar: 100 μm. **D.** Schematic of SPARCL1 treatment of BMDMs. **E.** Representative gating scheme for M2-like (F4/80^+^CD206^+^) macrophages after SPARCL1 treatment. **F.** Quantification of the proportion of M2-like (F4/80^+^CD206+) macrophages in M2 polarized BMDMs after treated with SPARCL1(10 μg/ml) for 24 hours. **G.** qPCR analysis for M2 macrophage genes (*Chil3* and *Mrc1*) in M1 and M2 polarized BMDMs after treated with SPARCL1(10 μg/ml) for 24 hours, n = 3 per group. **H.** BMDMs were pre-treated with TLR4 inhibitor, TAK-242(10 μM) for 1 hour and then incubated with or without SPARCL1 (10 μg/ml) for 1 hour. Phosphorylation of NF-κB was detected by western blot assay, n = 3 experiments. **I.** BMDMs were pre-treated with TLR4 inhibitor, TAK-242(10 μM) for 1 hour and then incubated with or without SPARCL1 (10 μg/ml) for 24 hours, and cell supernatant was collected. Th1 cytokines IL-1β, IL-6 and TNF-α levels in supernatant were measured by ELISA, n = 3 per group. **J.** GSEA analysis indicates that endothelial overexpression of Sparcl1 *in vivo* positively engaged the TLR4 signaling in isolated macrophages (see Fig. 4E-G). **K.** The TLR4 inhibitor TAK-242 blocks SPARCL1-induced NF-κB activation, thereby inhibiting macrophage transition to a pro-inflammatory phenotype. Data in (**B**) and (**I**) are presented as means ± SEM, calculated using one-way analysis of variance (ANOVA), followed by Dunnett’s multiple comparison test. Data in (**G**) are presented as means ± SEM, calculated using unpaired two-tailed t test; *P < 0.05, **P < 0.01, ***P < 0.001.

SPARCL1 has been reported to directly bind Toll-like receptor 4 (TLR4) in hepatocytes to activate a downstream inflammatory response cascade(24), and stimulation of TLR4 directly activates NF-κB (35). We therefore utilized the TLR4 specific inhibitor, TAK-242 (10 μM) to treat BMDMs 1 hour prior to SPARCL1 treatment (10 μg/ml). TAK-242 almost completely blocked SPARCL1-induced phosphorylation of NF-κB p65 (**Fig.5H**) and downstream release of TNFα, IL-1β and IL-6 (**Fig.5I**). GSEA revealed EC overexpression of Sparcl1 positively correlated with the TLR4 receptor pathway, and we observed upregulation of TLR4 downstream genes in EC^Sparcl1-OE^ mouse macrophages (**Fig.5J** and **Suppl Fig.7E**). Taken together, these data strongly indicate that SPARCL1 induces an M1 proinflammatory macrophage phenotype by activating TLR4/NF-κB signaling (**Fig.5K**).

### TAK-242 administration ameliorates pneumonia exacerbations induced by endothelial SPARCL1 overexpression

Having established that EC overexpression of Sparcl1 worsens viral pneumonia through TLR4 signaling, we next explored whether inhibition of TLR4 is sufficient to ameliorate influenza-induced injury in mice. To test this hypothesis, WT and EC^Sparcl1-OE^ mice were treated with TAK-242(3 mg/kg, i.p, every 2 days) from day 7 to 20 post influenza infection and mouse lungs were harvested on day 20 or 25 (**Fig.6A**). TAK-242 treatment significantly improved pneumonia symptoms in EC^Sparcl1-OE^ mice as assessed by reduced weight loss, shorter recovery time (**Fig.6B** and **Suppl Fig.8A**), and improved gas exchange as indicated by higher oxygen saturation (**Fig.6C** and **Suppl Fig.8B**) compared with vehicle treatment. Moreover, TAK-242 treatment significantly reduced M1_like and slightly increased M2_like macrophages (**Fig.6D-F**) and reduced pro-inflammatory cytokines levels (**Fig.6G**) including TNFα, IL-1β and IL-6 in EC^Sparcl1-OE^ mice on day 20 after infection. Intriguingly, TAK-242 did not exhibit appreciable therapeutic effects in WT mice, including no significant differences in body weight loss (**Suppl Fig.8A**), oxygen saturation levels (**Suppl Fig.8B**), changes in M1/M2 macrophage populations (**Suppl Fig.8C and D**) and BALF cytokines (with the exception of TNFα) (**Fig. 6G**). We interpret these observations to indicate that some degree of TLR4-mediated inflammatory signaling is beneficial, therapeutic targeting of TLR4 may require a strict administration time window, and inhibition of TLR4 may only be therapeutic in individuals with very high levels of SPARCL1. Taken together, targeting TLR4 signaling effectively alleviates the exacerbation of inflammation caused by overexpression of SPARCL1 in ECs, confirming that SPARCL1 mediates downstream inflammatory responses through TLR4 signaling *in vivo.* These findings also spur the prediction that if patients exhibit heterogeneity in SPARCL1 levels upon viral lung injury, treatment with TLR4 / NF-kB inhibitors may be therapeutic for a subset of these patients with the highest SPARCL1 levels.

**Fig.6.**
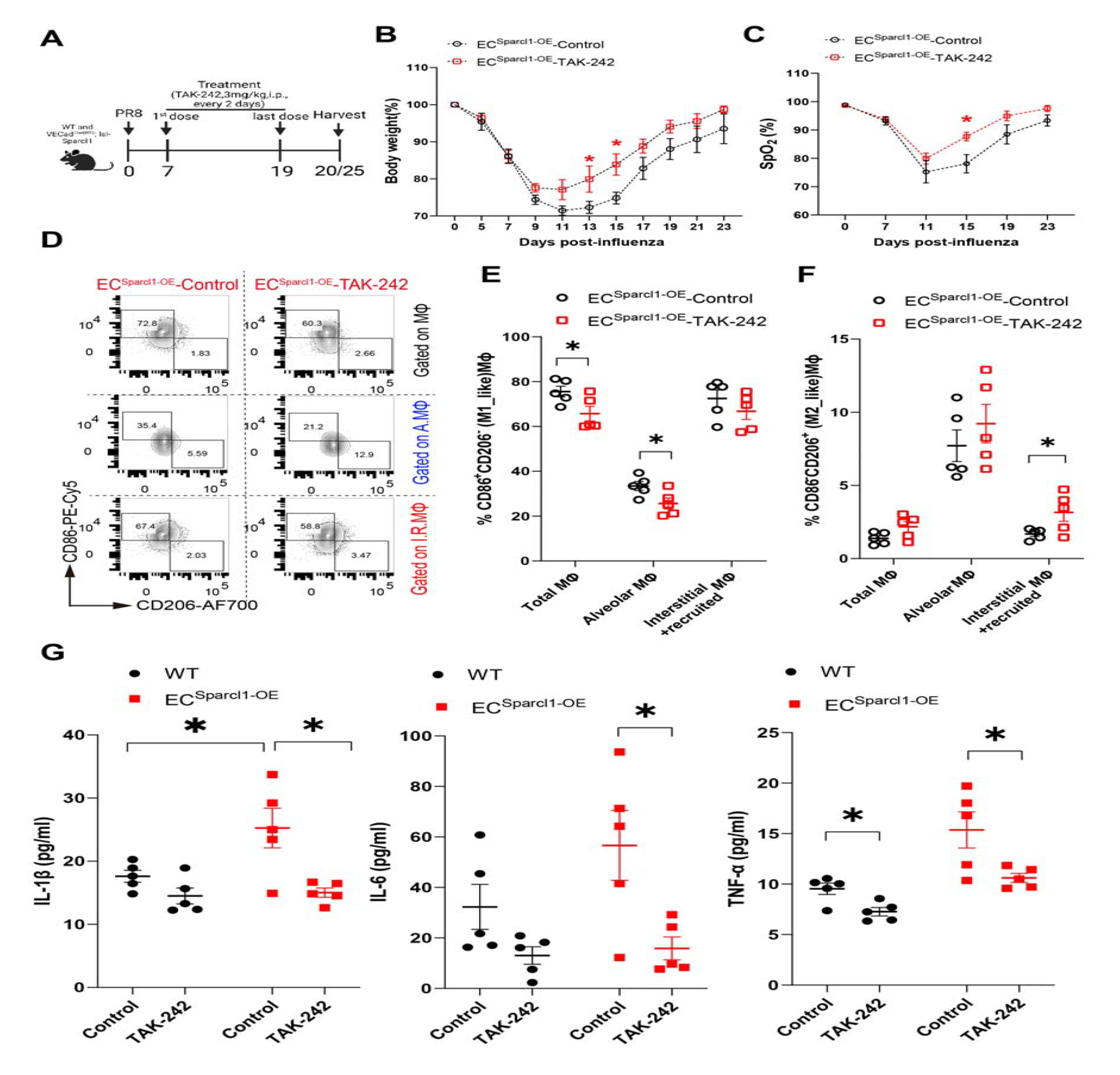
Blockade of TLR4 ameliorates the exacerbation of pneumonia induced by endothelial overexpression of Sparcl1. **A.** Timeline for TAK-242 administration and sampling. WT and EC mice were treated with TAK-242 (3 mg/kg, i.p.) or equal volume of vehicle control (DMSO). **B-C.** Time course of changes in (**B**) body weight and (**C**) capillary oxygen saturation in EC^Sparcl1-OE^ mice treated with or without TAK-242 after influenza infection, n = 4-7 mice per group. **D**. Representative gating scheme for identification of M1-like (CD86^+^CD206^-^) and M2-like (CD86^-^CD206^+^) macrophages in total lung macrophages (CD45^+^Ly6G^-^CD64^+^F4/80^+^), alveolar macrophages (CD45^+^Ly6G^-^CD64^+^F4/80^+^SiglecF^+^), and interstitial and recruited macrophages (CD45^+^Ly6G^-^CD64^+^F4/80^+^SiglecF^-^) at day 20 after influenza infection in EC^Sparcl1-OE^ mice treated with or without TAK-242. **E-F**. Quantification of the proportion of (**E**)M1-like (CD86^+^CD206^-^) and (**F**)M2-like (CD86^-^ CD206^+^) macrophages in total lung macrophages (CD45^+^Ly6G^-^CD64^+^F4/80^+^), alveolar macrophages (CD45^+^Ly6G^-^CD64^+^F4/80^+^SiglecF^+^) and interstitial and recruited macrophages (CD45^+^Ly6G^-^CD64^+^F4/80^+^SiglecF^-^) at day 20 after influenza infection in WT and EC^Sparcl1-OE^ mice treated with or without TAK-242, n = 5 mice per group. **G**. The levels of Th1 cytokines, IL-1β, IL-6 and TNF-α in bronchoalveolar lavage fluid (BALF) were measured by ELISA in WT and EC^Sparcl1-OE^ mice treated with or without TAK-242 at day 20 after influenza infection, n = 5 mice per group. Data in (**B**), (**C**), (**E**) and (**F**) are presented as means ± SEM, calculated using unpaired two-tailed t test; Data in (**G**) are presented as means ± SEM, calculated using one-way analysis of variance (ANOVA), followed by Dunnett’s multiple comparison test. *P< 0.05.

### High levels of SPARCL1 are associated with poor outcomes in patients with viral pneumonia

To determine the clinical relevance of our animal-based observations, lungs from COVID- 19 ARDS patients (collected after viral clearance, COVID_donors) and healthy control lungs (Healthy_donors) were collected. Immunostaining for SPARCL1 revealed higher expression of SPARCL1 in vascular ECs (ERG^+^SPARCL1^+^) in post-COVID lungs compared to healthy lungs (**Fig.7A**). qPCR analysis of ECs isolated from post-COVID and healthy lungs indicated that *SPARCL1* was significantly upregulated in COVID endothelium (**Fig.7B**). Survival analysis demonstrated that the level of SPARCL1 in the plasma of patients with fatal COVID-19 disease was significantly higher than that of those who survived (**Fig.7C**). These observations reinforce our findings *in vitro* and in our transgenic mouse models, indicating that high levels of SPARCL1 are closely related to the development and exacerbation of COVID-19 pneumonia, and detection of SPARCL1 in plasma represents a potential method to evaluate the prognosis of viral pneumonia as well as to potentially identify patients who may response positively to SPARCL1, TLR4, or NF-κB inhibition.

**Fig.7.**
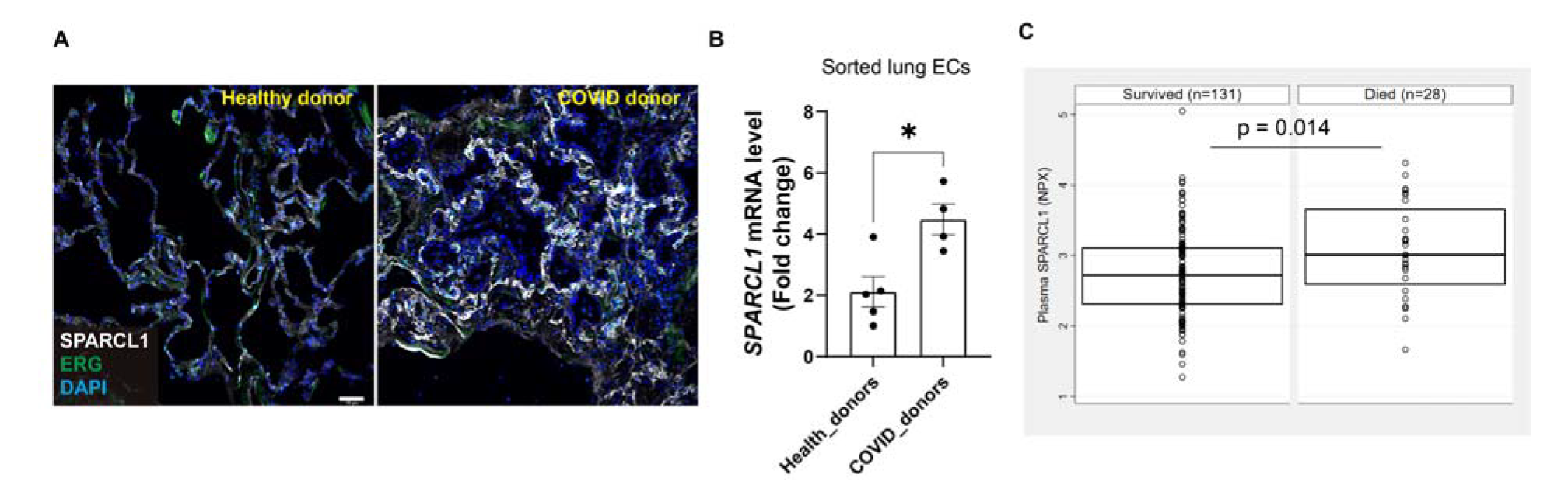
High expression of SPARCL1 in the pulmonary vascular endothelium of COVID-19 patients correlates with increased mortality. **A.** Representative immunostaining image of SPARCL1 in endothelial cells (ERG) in both healthy and COVID-19 donors’ lung tissue. Scale bar, 50 μm. **B.** qPCR analysis of *SPARCL1* in isolated lung ECs (CD45^-^EpCAM^-^CD31^+^) sorted from both healthy and COVID-19 donors’ lung tissue, n = 4-5 per group. **C.** The SPARCL1 level in plasma from COVID-19 patients was measured by Olink proximal extension assay. Data are presented as means ± SEM, Data in (B) were calculated using unpaired two-tailed t test; Data in (C) were calculated using Wilcoxon rank sum test, *P< 0.05.

## Discussion

Endothelial cells (ECs) lining the capillary network around the alveoli play a crucial role in lung physiology, responsible for gas exchange, nutrient transport, and leukocyte trafficking. Although they are not typically directly infected by respiratory viruses, viral infections can nonetheless trigger a severe local inflammatory response leading to EC apoptosis, necrosis, and other forms of cell death(8). ECs undergo a continuous and dynamic adjustment of their functions in response to pathogens / damage-associated molecular patterns or cytokines, leading to the classical “endothelial activation” phenotype characterized by induction of adhesion molecules ICAM-1, VCAM-1, and E-selectin(36). This leads to an increased adherence of leukocytes to the endothelial surface and heightened transendothelial migration, facilitating influx of immune cells which, in turn, contribute to both injury and repair processes in the tissue. As in other organs, lung ECs are heterogeneous, consisting of arterial, venous, lymphatic, and two distinct subsets of capillary ECs, aerocytes (aCap) ECs and general capillary (gCap) ECs, thought to govern gas exchange and capillary repair, respectively(26, 28). While various studies have investigated the differences in the responses of lung EC types to injury and found evidence of functional differences among these subpopulations, the lack of strict and reliable tracing markers has limited their conclusions, which are primarily based on bioinformatic prediction. Further experimental evidence is needed to fully validate the functional consequences of these transcriptomic differences.

In this study, using single cell transcriptome analysis, we uncovered two "active" subpopulations of gCap ECs that exhibited alternating behavior during influenza-induced lung injury. By analyzing their characteristic genes, we found that they were similar to the two subpopulations of ECs recently described by Zhang et al (27), so we adopted the naming conventions of Dev.ECs and Immu.ECs. Of note, we observed dramatically reduced numbers of Dev.ECs on day 20 post-influenza, but by day 30, when lung respiration and oxygen saturation had largely returned to normal, Dev.ECs had been largely restored, indicating these cells likely contribute to lung homeostasis. Further analysis revealed that the Dev.ECs highly expressed *Atf3*, a known stress-induced transcriptional regulator (37), suggesting that endothelial *Atf3* may be involved in maintaining lung homeostasis, reinforced by a recent study demonstrating the importance of Atf3-expressing ECs in lung regeneration (38). The Immu.ECs, which primarily appear after injury and may be unique to lung injury / repair, have received limited attention in previous studies. Thus, we focused on these cells. Further analysis revealed that the Immu.ECs highly expressed *Sparcl1*, and given the paucity of functional studies on this gene, we pursued further characterization.

SPARCL1, a matricellular protein and member of the SPARC protein family, has been described as a blood vessel–derived anti-angiogenic (“angiostatic”) protein(25). However, overexpression or deletion of SPARCL1 in ECs *in vivo* did not show obvious effects on EC proliferation. SPARCL1 is also expressed in pericytes / mural cells in some tissues and is reported to be required for vascular maturation and integrity(25, 39). In the lung, however, capillary ECs (especially aerocytes) are largely discontinuous with pericytes (40), so we postulated that SPARCL1 expression from pericytes may be less relevant in this context. Instead of direct effects on the vasculature, our findings suggested that the high expression of SPARCL1 by ECs during pneumonia contributes to the worsening of lung injury by driving Th1 inflammation, ultimately causing more harm than benefit. Our ELISA data showed that SPARCL1 was significantly upregulated during pneumonia before gradually returning to normal levels during recovery, so we speculated that the high expression of SPARCL1 in influenza-induced lung injury was related to the level of local inflammation. These findings reinforce previously described studies showing that SPARCL1 activates the hepatic inflammatory response and contributes to liver injury in steatotic mice(24), and our work here points to macrophage activation as the major driver of this SPARCL1-mediated inflammatory exacerbation.

Macrophages play a pivotal role in orchestrating the initiation and resolution of inflammation, as well as repair responses in the lungs (41). The diversity of macrophage function is often “binned” into polarized states, with M1 and M2 subtypes implicated as both drivers and regulators of disease (32). It has been demonstrated that either persistence of inflammatory macrophage numbers or prevention of their conversion to a reparative anti-inflammatory phenotype can further delay tissue repair following injury (42). Therefore, identifying and characterizing the mechanisms that drive macrophages to exhibit pro or anti-inflammatory activity is critical to promote resolution of tissue inflammation/repair responses. We show vascular-derived SPARLC1 exacerbates pneumonia by agonizing TLR4 and promoting the polarization of pro-inflammatory macrophages. Further, we also observed that SPARCL1 converted M2-like reparative macrophages toward a more M1-like state. These results indicate that 1) macrophages are not permanently committed to a singular activation state, instead exhibiting the plasticity necessary to convert between polarized states, and that 2) the activation status of macrophages, as well as the inflammatory microenvironment wherein they reside, are crucial factors influencing their response to and function within injured lungs. Intriguingly, similar effects on macrophage activation in adipose tissue have been described for SPARC (43), suggesting the SPARC matricellular protein family may share a conserved function in modulating macrophage activity.

Though our work here elucidates novel mechanisms of endothelial-macrophage crosstalk and macrophage activation, important questions remain. While not formally demonstrated here, all indications are that Immu.ECs essentially represent an “activated / inflamed” state of gCaps rather than a *de novo* population. If true, the initiating signals driving the transcriptomic and phenotypic switch from Dev.ECs to Immu.ECs remain unknown. Although it has been demonstrated *in vitro* that *SPARCL1* expression can be induced by Th1 pro-inflammatory cytokines dependent upon cell confluency(25), it is difficult to know what exactly this confluency-dependent effect means *in vivo*, where the endothelium is already “confluent” at steady state. In addition, while our transgenic mouse models clearly indicate that the endothelial source of SPARCL1 is critical for the observed phenotypes, our work does not address whether there may be an additive role for mesenchymal sources of SPARCL1, nor does it assess whether SPARC might synergize with SPARCL1 to influence macrophage behavior in the lung.

While multiple studies have established the importance of resident and recruited lung macrophages in both pulmonary inflammation and tissue repair processes, much remains to be discovered regarding the underlying signaling pathways and mechanisms of multilineage paracrine communication involved. Our findings reveal, for the first time, that the vascular-derived molecule SPARCL1 is a critical paracrine signaling factor between capillary endothelial cells and macrophages, and that macrophage responses to SPARCL1 primarily contribute to the severity of pneumonia. In this sense, SPARCL1 should be considered a driver of pathological inflammation, and while SPARCL1 may have important tissue protective effects as well (44), our model points to a threshold beyond which SPARCL1 primarily drives overexuberant inflammation. Integration of our findings in mouse models and patient samples leads us to suggest that assessing SPARCL1 levels in pneumonia patients serves not only as a biomarker of disease severity, but may allow for personalized medicine approaches. In patients with particularly high SPARCL1 levels, treatment with SPARCL1 / TLR4 / NF-κB antagonists could blunt uncontrolled inflammation, help them weather the cytokine storm, and ultimately promote better outcomes for these patients.

## Methods and materials

### Patient samples

Human lung samples (both normal lung and COVID-19 samples) were obtained from Penn Lung Biology Institute Human Lung Tissue Bank (https://www.med.upenn.edu/lbi/htlb.html). COVID-19 samples were from patients who previously tested positive for COVID-19 by PCR but tested negative via PCR multiple times prior to tissue acquisition. All COVID-19 samples were obtained from ventilated ARDS patients at least 30 days post-hospitalization who underwent lung transplant, at which time tissue samples were acquired.

Human plasma samples were from a prospective cohort study of participants with or high risk for sepsis (**Table S2**). To be eligible, subjects were admitted to the hospital and tested positive for SARS-CoV-2 by PCR test of nasal or respiratory secretions as we have published (45, 46). Subjects were excluded if they had been previously enrolled to the cohort, if they were chronically critically ill and residing in a long term advanced care hospital, if they desired exclusively palliative care on admission, or if the participant or their proxy were unable or unwilling to consent to the study. Blood was sampled within 3 days of hospital admission, processed immediately for plasma, and frozen at −80°C until analysis. Plasma was not treated for viral inactivation prior to assay. Trained study personnel collected demographic and clinical data from the electronic health record (EHR) into case report forms. Subjects were characterized by the World Health Organization ordinal scale for respiratory failure (47) at the time of blood sampling and considered severe respiratory failure if they required high flow oxygen (>6 lpm), non-invasive ventilation, or invasive ventilation (WHO ordinal scale ≥ 6) and moderate respiratory injury if they required no oxygen or oxygen at flow rates at or below 6 lpm (WHO ordinal scale ≤ 5) (45). Mortality was assessed at 90 days using the EHR, which included a surveillance program post-discharge for patients discharged after COVID-19.

We assayed plasma proteins with the Olink proximal extension assay as described (45) and filtered results for SPARCL1. Protein concentrations were compared between categorical groups using the Wilcoxon rank sum test.

### Generation of Sparcl1^flox^ mice

Sparcl1^flox^ mice were generated using CRISPR-Cas9 strategy in mouse embryonic stem cells (ESCs). Briefly, DNA template containing loxP sites flanking exon 2 of the Sparcl1 “loxP-Sparcl1_Ex2-loxP” was synthesized by Genscript (Genscript Biotech Corp.) and purified. The gRNAs targeting the flanks of Sparcl1 exon2 were cloned into a CRISPR-Cas9 vector, pDG459 (Addgene, #100901) according to the protocol described previously with minor modifications(48). Mouse ESCs(129/B6 F1 hybrid ES cell line V6.5)(49) were electroporated with linearized donor DNA (loxP-Sparcl1_Ex2-loxP) and pDG-459 with 2 gRNAs inserted and clones were selected by puromycin (2ug/ml) for 4-5 days. Following successful homologous recombination between the targeting donor DNA and ESC cell DNA as judged by PCR screening, targeted ESCs were then injected into C57BL/6J blastocysts to obtain chimeric mice following standard procedures. Chimeric mice (Sparcl1^flox/+^) are bred with VECad^CreERT2^ (Cdh5^CreERT2^) female mice (50) for direct characterization. Genotyping primers and PCR program were listed in **table S1**.

### Generation of Sparcl1^OE^ (R26-LSL-Sparcl1) mice

Mouse Sparcl1 cDNA was amplified by PCR using the following primers: forward primers with 5’ arm Xho I restrict enzyme cutting site added, mSparcl1-XhoI-F: ccgctcgagcggatgaaggctgtgctt ctcctc, reverse primers with 5’ arm Sac II restrict enzyme cutting site added, mSparc1-SacII-R: tccccgcggggatcaaaagaggaggttttcatctat. Sparcl1 PCR products were cut, purified, and inserted into a generic targeting vector (pBigT, Addgene, #21270), pBigT-mSparcl1 was then subsequently cloned into a plasmid with the ROSA26 genomic flanking arms (ROSA26-PA, Addgene, #21271) following the protocol described previously (51), generating the final targeting vector ROSA-26-pBigT-mSparcl1 for homologous recombination. The linearized targeting vector was electroporated into mouse ESCs (same line as described above), and G418-resistant colonies were analyzed for proper editing by PCR. Finally, the targeted ESCs were injected into C57BL/6J blastocysts, and the resulting chimeric mice were bred with VECad^CreERT2^ (Cdh5^CreERT2^) female mice for direct characterization. Genotyping primers and PCR program were listed in **table S1**.

### Animal treatments

Sparcl1^flox^ (noted as Sparcl1^flox/flox^) or R26-LSL-Sparcl1 (noted as Sparcl1^+/WT^ or Sparcl1^+/+^) mice were crossed with VECad^CreERT2^ (Cdh5^CreERT2^) mice (50) to produce VECad^CreERT2^; Sparcl1^flox/flox^ mice, VECad^CreERT2^; Sparcl1^+/WT^, and VECad^CreERT2^; Sparcl1^+/+^ mice. Sparcl1^flox/flox^ mice lacking Cre or VECad^CreERT2^; Sparcl1^WT/WT^ mice were used as the conditional endothelial Sparcl1 knockout or overexpression control mice, respectively; These mice were administered five doses of tamoxifen (0.25 mg/g body weight) in 50 μl of corn oil every other day and rested for 2 weeks after the last injection, resulting in EC-specific deletion (EC^Sparcl1-KO^) or overexpression (EC^Sparcl1-OE^) of Sparcl1 in adult mice. Afterward, influenza virus A/H1N1/PR/8 was administered intranasally at 50 to 75 TCID50 units to mice according to experimental requirements as our previously described(9, 52). Mice were weighed regularly and euthanized at the indicated time points for tissue harvest. In this study, all mice were used at 6 to 8 weeks old, and mice of both sexes were used in equal proportions. All animal experiments were carried out under the guidelines set by the University of Pennsylvania’s Institutional Animal Care and Use Committees and followed all National Institutes of Health (NIH) Office of Laboratory Animal Welfare regulations.

### Cell culture conditions

Bone marrow-derived macrophages (BMDMs) were prepared as described previously(53). Briefly, bone marrow cells were collected from the femur and tibia of C57BL/6 mice and cultured in RPMI 1640 media with GlutaMAX supplement added, containing 10% cosmic calf serum (CC; HyClone, #SH3008704), 1% penicillin/streptomycin (P/S; Gibco, #15140122) and 20 ng/ml recombinant murine M-CSF (PeproTech, #315-02). Depending on the experiment, BMDMs were treated with recombinant mouse SPARCL1 (5-20 μg/ml, R&D system, #4547-SL; SinoBiological, # 50544-M08H); recombinant IL-4 (20 ng/ml, PeproTech, #214-14), lipopolysaccharides (50 ng/ml; Sigma Aldrich, #L6529), Resatorvid (TAK-242) (10μM; MedChemExpress, # HY-11109) or vehicle control.

### BALF cell count and total protein quantification

Bronchoalveolar lavage fluid (BALF) was collected by inserting a catheter in the trachea of euthanized mice. Lungs were infused with 1 ml PBS and gently retracted to maximize BALF retrieval and to minimize shear forces. The fluid was centrifuged for 5 min at 500×g, the supernatant was collected for downstream experiments such as total protein quantification and ELISA, cell pellets were re-suspended and red blood cells were removed using Red Blood Cell Lysis Buffer (Thermo Fisher Scientific, A1049201). The total cell count was determined using a cell counting chamber under light microscopy. Total protein in BALF was determined by the bicinchoninic acid (BCA) colorimetric assay using Pierce BCA Protein Assay Kit (Thermo Fisher Scientific, #23227).

### ELISA

BALF was collected in 1 ml PBS as mentioned above. SPARCL1 (Mybiosource, #MBS2533471), TNF-α (Invitrogen, #88-7324-22), IL-6 (Invitrogen, #88-7064-22), and IL-1β (Invitrogen, #88-7013-22) levels in mouse BALF were measured by ELISA assessments according to the manufacturer’s instructions.

### Whole lung cell suspension preparation

Human lung single-cell suspensions were prepared as described previously with slight modifications (54). Briefly, distal lung tissue was obtained and dissected into roughly 5□cm^3^ pieces. Tissue was washed in 200□ml sterile PBS for 5□min at 4□°C at least two times, or until PBS no longer appeared obviously bloody. An additional 5□min wash was then performed with Hank’s buffered saline solution (HBSS). Using autoclaved Kim Wipes, tissue was compressed to remove as much liquid as possible and further dissected into<1□cm^3^ pieces. Sterile HBSS buffer containing 5□U/ml Dispase II and 0.1□mg/ml DNase I + penicillin/streptomycin was added to the small tissue pieces. Tissue rapidly takes up the digest solution at this point, becoming visibly engorged. Tissue was digested on a shaker at 220 rpm at 37°C for 2□hours. Tissue was then liquified in digestion solution using an Osterizer Blender as follows: (low setting for all) 5□s milkshake, 3□s smoothie, and 5□s milkshake. The suspension was poured through a glass funnel lined with sterile 4 × 4 gauze, applying some compression to recover as much of the solution as possible. The cell suspension was sequentially filtered through 100□μm, 70□μm and 40□μm strainers (Thermo Fisher Scientific). Finally, red blood cells were removed using Red Blood Cell Lysis Buffer (Thermo Fisher Scientific, A1049201).

Lungs were harvested from mice and single-cell suspensions were prepared as previously described(9). Briefly, the lungs were thoroughly perfused with cold PBS via the left atrium to remove residual blood in the vasculature. Lung lobes were separated, collected, and digested with collagenase II (5 mg/ml in HBSS) (Worthington Biochemical, #LS004176) for 1 hour at 37 □ on shaker at speed of 200 rpm, and mechanically dissociated by pipetting in sort buffer (DMEM + 2% CC + 1% P/S; referred to as “SB”). Next, cell suspensions were filtered by the 70-μm cell and treated by red blood cell lysis buffer containing 1:500 deoxyribonuclease I (DNase I) (MilliporeSigma, #D4527) for 5 min at room temperature, and the cell suspension was then used for subsequent experiments.

### Fluorescence-activated cell sorting and analysis

Whole lung single-cell suspensions were prepared as above; BMDMs were treated according to the experimental requirements, and Accutase (Sigma Aldrich, #A6964) was used to digest into single-cell suspension. Single-cell suspension were then blocked in SB containing 1:50 human or mouse TruStain FcX™ for 5-10 min at 37°C. The cell suspension was stained using cell viability dye (1:1000 in PBS, eBioscience^TM^, #65-0865-18, intracellular FACS analysis use only) allophycocyanin (APC)/Cyanine7 or Brilliant Violet 421™ anti-human CD45 antibody (1:200, Biolegend, HI30), APC anti-human CD31 antibody (1:200, Biolegend, WM59) and PE anti-human CD326 (EpCAM) antibody (1:200, Biolegend, 9C4) for human lungs; Brilliant Violet 785™ anti-mouse CD45 antibody (1:200; Biolegend, 30-F11), Brilliant Violet 711™ anti-mouse/human CD11b antibody (1:200; Biolegend, M1/70), Brilliant Violet 421™ anti-mouse F4/80 antibody(1:100; Biolegend, BM8), Alexa Fluor® 647 anti-mouse Siglec-F antibody (1:100; BD Bioscience, E50-2440), PE anti-mouse Ly-6G antibody(1:200; Biolegend, 1A8), PE/Cyanine7 anti-mouse CD64 (FcγRI) antibody(1:200, Biolegend, X54-5/7.1), Alexa Fluor 488 or PE–conjugated rat anti-mouse CD31 [platelet endothelial cell adhesion molecule 1 (PECAM1)] antibody (1:200; BioLegend, MEC13.3), Alexa Fluor 647 or FITC–conjugated rat anti-mouse CD326 (Ep-CAM) antibody(1:200; Biolegend, G8.8), PE/Cyanine5 anti-mouse CD86 antibody(1:200; Biolegend, GL-1), PE/Cyanine5 anti-mouse CD3ε antibody (1:200; Biolegend, 145-2C11), Alexa Fluor 700 anti-mouse/human CD45R/B220 antibody(1:50; Biolegend, RA3-6B2), BUV395 anti-mouse CD11b(1:200; BD Biosciences, M1/70), Brilliant Violet 421™ anti-mouse NK-1.1 antibody(1:100; Biolegend, PK136), PE-Cyanine7 CD127 Monoclonal antibody (1:100; Invitrogen, A7R34) for 45 min at 4°C. Stained cells and “fluorescence minus one” (FMO) controls were then resuspended in SB + 1:1000 DNase + 1:1000 Draq7 (BioLegend, #424001) as a live/dead stain. All flow analyses were performed on BD FACSymphony A3 Cell Analyzer (BD Biosciences) and FACS sorting was performed on a BD FACSAria Fusion Sorter (BD Biosciences).

### Intracellular FACS analysis

Cell surface antigen antibodies were prepared and stained as describe above, and then single-cell suspensions were fixed and permeabilized using Cyto-Fast™ Fix/Perm Buffer Set (Biolegend, #426803) according to the manufacturer’s instructions. The cells were then stained with Alexa Fluor® 700 anti-mouse CD206 (MMR) antibody (1:200; Biolegend, C068C2) for 30 min at room temperature; For intracellular EdU cytometry flow, mice were injected intraperitoneally with EdU (50 mg/kg; Santa Cruz Biotechnology, #sc-284628) at indicated time points. After euthanasia, the whole lung single-cell suspension was prepared and stained as above and fixed by 3.2% paraformaldehyde (PFA) (Electron Microscopy Sciences, #15714-S) for 15 min, washed twice using 3% bovine serum albumin (BSA) (in PBS), and permeabilized using 0.1% Triton X-100 (in PBS) for 15 min. EdU was detected using the Click-iT reaction coupled to an Alexa Fluor 647 azide following the instructions of the manufacturer (Invitrogen, #C10086). Intracellular flow analyses were performed on BD FACSymphony A3 Cell Analyzer (BD Biosciences).

### Immunofluorescence

For cryostat tissue sections, human lungs were obtained and transported to the laboratory on ice and mouse lungs were isolated and processed as previously described(55). Freshly dissected lungs were fixed, embedded and cut into 7 μm thick cryosections, and postfixed another 5 min with 3.2% PFA. Tissue sections were blocked in blocking buffer (1% BSA, 5% donkey serum, 0.1% Triton X-100, and 0.02% sodium azide in PBS) for 1□hour at room temperature. Afterward, slides were probed with primary antibodies (CD31 1:200, Biolegend, MEC13.3; mSPARCL1 1:500, R&D systems, #AF2836-SP; hSPARCL1 1:500, R&D systems, #AF2728-SP; ERG 1:2000, Abcam, #ab92513; F4/80 1:200, Cell Signaling Technology, #30325; RELMα 1:200; Invitrogen, #56-5441-82) and incubated overnight at 4°C. The next day, slides were washed and incubated with the fluorophore-conjugated secondary antibodies (typically Alexa Fluor conjugates, Life Sciences) at a 1:1000 dilution for ≥1 hour. Last, slides were again washed, incubated with 1 μM 4′,6-diamidino-2-phenylindole (DAPI) for 5 min, and mounted using ProLong Gold (Life Sciences, #P36930). 4-6 images were taken randomly from each sample/section with a Leica Dmi8 microscope and analyzed with LAS X software (Leica).

### Histological analysis

Lung tissue sections fixed with 3.2% PFA were stained with Hematoxylin and Eosin Stain Kit (Vector Laboratories, #H-3502) according to the manufacture’s instruction and then imaged with a Leica DMi8 microscope.

### Pulse oximetry

Repeated measurements of peripheral oxygen saturation (SpO2) were taken using a MouseOx Plus rat & mouse pulse oximeter and a MouseOx small collar sensor (Starr Life Sciences Corp.). Mice were shaved around the neck and shoulders where the collar sensor sits. Recordings were taken using MouseOx Premium Software (Starr Life Sciences Corp., Oakmont, PA, USA). Measurements were taken continuously for >3□min at a measurement rate of 15 Hz. Measurements were imported into Microsoft Excel, and all readings with a nonzero error code were filtered out. The average of these error-free readings was used to calculate the SpO2 reading for each mouse for each given time point.

### Western blotting

Total protein from cells was extracted by lysis in radioimmunoprecipitation assay buffer (Santa Cruz Biotechnology, #sc-24948) with protease inhibitor cocktail (Cell Signaling Technology, #5872). Protein concentrations were determined using a BCA protein assay kit (Thermo Fisher Scientific, #23227). Samples with equal amounts of protein were fractionated on SDS–polyacrylamide gels (Bio-Rad, #4561084), transferred to polyvinylidene difluoride (MilliporeSigma, #IPVH00005) membranes, and blocked in 5% skim milk (Cell Signaling Technology, #9999s) in TBST (0.1% Tween 20 in tris-buffered saline) for 1.5□hours at room temperature. The membranes were then incubated at 4°C overnight with primary antibodies [phospho-NF-κB p65 1:1000, Cell Signaling Technology, #3033; NF-κB p65 1:1000, Cell Signaling Technology, #8242; mSPARCL1 1:500, R&D systems, #AF2836-SP; β-actin 1:2000, Cell Signaling Technology, #4970]. After the membranes were washed with TBST, incubations with 1:4000 dilutions (v/v) of the secondary antibodies were conducted for 2□hours at room temperature. Protein expression was detected using the ChemiDoc XRS+ System (Bio-Rad). β- Actin was used as a loading control.

### RNA isolation and qPCR

Total RNA was extracted by ReliaPrep RNA Cell Miniprep kit according to the manufacturer’s recommendation (Promega, #Z6011) and then reverse-transcribed into complementary DNA using the iScript Reverse Transcription Supermix (Bio-Rad, #1708841). qPCR was performed using a PowerUp SYBR Green Master Mix and standard protocols on an Applied Biosystems QuantStudio 6 Real-Time PCR System (Thermo Fisher Scientific). Glyceraldehyde-3-phosphate dehydrogenase (GAPDH) was used to normalize RNA isolated from human ECs; RPL19 was used to normalize RNA isolated from mouse samples. The 2^−ΔΔCt^ comparative method was used to analyze expression levels. The primers used are listed in **table S1**. For sorted mouse lung ECs, RNA extracted and amplified (10 ng RNA/sample) using SMART-Seq® HT kit (Takara Bio, #634455) according to the manufacture’s instruction, the amplified cDNA was subsequently used for qPCR.

### Single-cell transcriptomics analysis

The whole lung single-cell suspension from mice on D0, D20 and D30 post influenza infection was prepared as above and FACS-sorted ECs were used for sequencing. Single-cell sequencing was performed on a 10X Chromium instrument (10X Genomics) at the Children’s Hospital of Philadelphia Center for Applied Genomics. Cellranger mkfastq was used to generate demultiplexed FASTQ files from the raw sequencing data. Next, Cellranger count was used to align sequencing reads to the mouse reference genome (GRCm38) and generate single cell gene barcode matrices. Post processing and secondary analysis was performed using the Seurat package (v.4.0). First, variable features across single cells in the dataset were identified by mean expression and dispersion. Identify variable features was then be used to perform a PCA. The dimensionally-reduced data was used to cluster cells and visualize using a UMAP plot. *Sparcl1* expression level was compared using VlnPlot package in Seurat. Sequencing data is available on GEO, https://www.ncbi.nlm.nih.gov/geo/query/acc.cgi?acc=GSE201631. Enter token uvidmokidfalvad.

### Bulk RNA-seq analysis

Whole lung single-cell suspensions from EC^Sparcl1-OE^ and WT mice on D20 post influenza infection were prepared as above and FACS-sorted macrophages (CD45^+^/Ly6G^-^/CD64^+^/F4/80^+^) were used for sequencing. RNA extracted and amplified (10 ng RNA/sample) using SMART-Seq® HT kit (Takara Bio, #634455) according to the manufacture’s instruction, the amplified cDNA was quality checked, DNA libraries were prepared, and sequencing was performed on an Illumina HiSeq platform by GENEWIZ Co. Ltd. Raw data (raw reads) in fastq.gz format were processed through a general pipeline as describe previously(56). Reads were aligned to the mm10 mouse genome using Kallisto and imported into R Studio for analysis via the TxImport package. Data was then normalized using the trimmed mean of M values normalization method in the EdgeR package. Mean-variance trend fitting, linear modeling, and Bayesian statistics for differential gene expression analysis were performed using the Voom, LmFit, and eBayes functions, respectively, of the Limma package, yielding differentially expressed genes between WT and EC^Sparcl1-OE^ groups. Based on PCA analyses, a single WT sample was a clear outlier from all other samples in both groups, likely due to poor sort purity, and was removed from subsequent analysis. Volcano plots were created using the OmicStudio tools at https://www.omicstudio.cn/tool. All detectable genes derived from RNA-seq were used for gene set enrichment analysis (GSEA) using the Molecular Signatures Database (MSigDB) C2: curated gene sets according to the standard GSEA user guide (http://www.broadinstitute.org/gsea/doc/GSEAUserGuideFrame.html). Sequencing data is available on GEO, https://www.ncbi.nlm.nih.gov/geo/query/acc.cgi?acc=GSE225439. Enter token cnwviccmzvodrmp.

### Statistics

All statistical calculations were performed using GraphPad Prism 9. All *in vitro* experiments were repeated at least 3 times unless otherwise stated. Unpaired two-tailed Student’s t-tests were used to ascertain statistical significance between two groups. One-way analysis of variance (ANOVA) was used to assess statistical significance between three or more groups with one experimental parameter. For details on statistical analyses, tests used, size of n, definition of significance, and summaries of statistical outputs, see corresponding figure legend and the Results section.

## Acknowledgements

We are grateful to the patients and families who agreed to participate and to the hospital staff who cared for them and facilitated our cohort study. We thank the Penn Lung Biology Institute Human Lung Tissue Bank, CHOP Flow Cytometry Core and The Penn Vet Imaging Core (PVIC) for their assistance in performing these studies. We thank Dr. Noam Cohen for facilitating additional access to flow sorters. Finally, we thank BioRender for providing a platform to create the cartoons and schematics used in figures throughout this report. This work was supported by the following funding sources: NIH R01HL153539 (AEV), NIH R01HL164350 (AEV), NIH R01HL161196 (NJM), NIH R01HL155804 (NJM), R01HL155821 (EC).

## Author contributions

Conception and design: G.Z., N.J.M, M.J.M., A.E.V.

Data acquisition: G.Z., L.X., A.I.W., M.E.G., C.V.C., S.A., J.W., X. L., S.K., N.P.H., M.C.B., K.M.S., J.D.P., E.C., J.D.C., M.M.C.

Data analysis: G.Z., N.J.M, and A.E.V.

Writing: G.Z., N.J.M., A.E.V.

All authors read and approved the final manuscript.

