## Supplementary Figures for "Vascular Endothelial-derived SPARCL1 Exacerbates Viral Pneumonia Through Pro-Inflammatory Macrophage Activation"

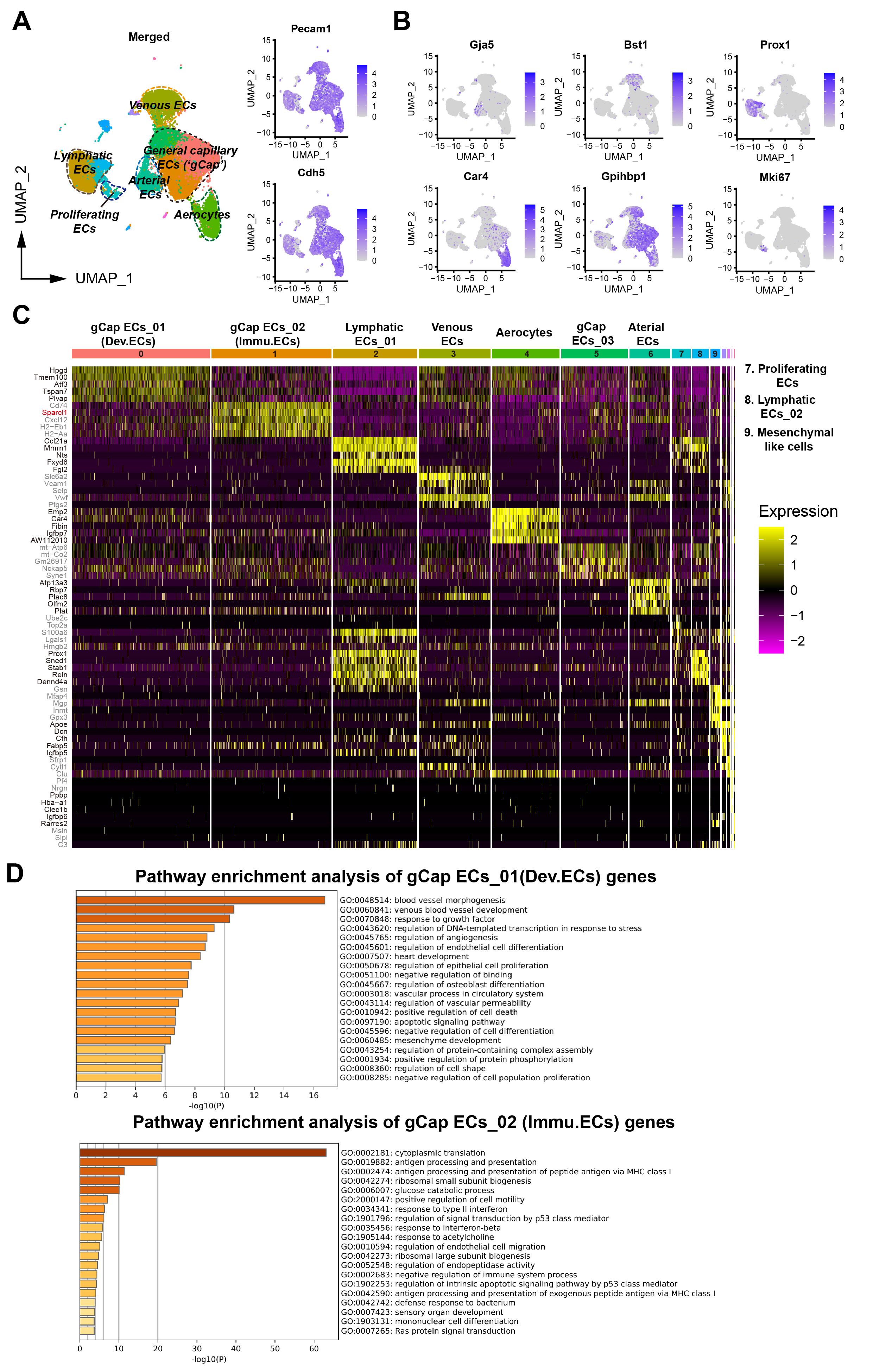
**Suppl Fig.1**

**Suppl Fig.1 Single-cell transcriptomics reveals heterogeneous changes in pulmonary vascular endothelial cells after influenza injury**

**A**. UMAP plots showed the signature EC genes *Pecam1*(CD31), *Cdh5* and the major EC clusters annotated for mouse lungs.

**B.** Feature plots for identified EC cluster signature genes(*1, 2*).

**C**. Heatmap of the most differentially expressed genes (Top 5) of lung ECs in all clusters. The color bars indicate gene expression level in log2 scale.

**D**. The GO terms of differentially expressed genes in Dev.ECs and Immu.ECs were carried out by Metascape(*3*).

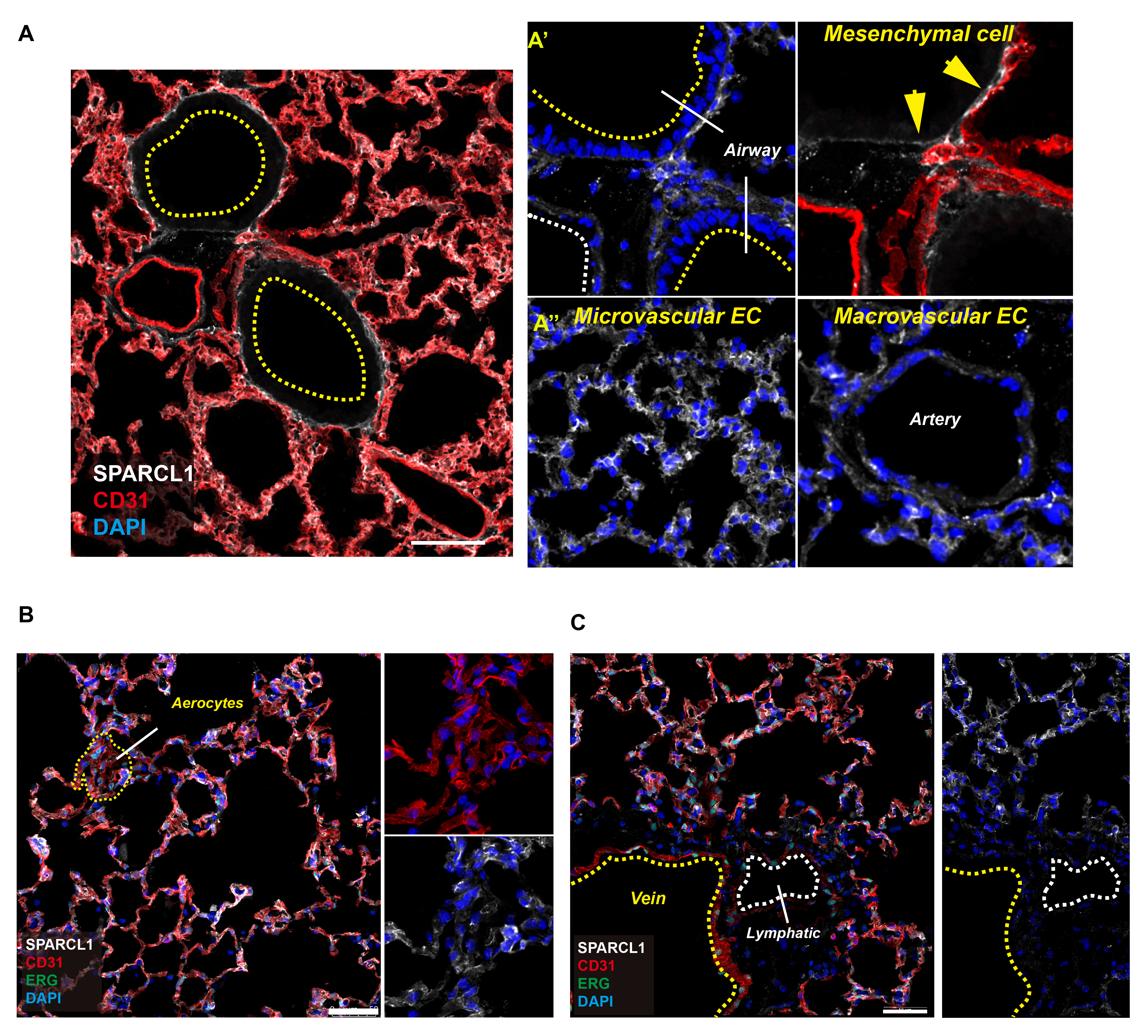
**Suppl Fig.2**

**Suppl Fig.2 In mouse lung, SPARCL1 is mainly expressed in capillary ECs, especially in gCap ECs**

**A**. Mouse lung tissues (uninjured) were stained with SPARCL1, ERG and CD31/PECAM1, demonstrating SPARCL1 expressed in stromal/mesenchymal cells (**A’**), lung capillary ECs (**A’’**), scale bar: 50 μm.

**B**. SPARCL1 is low / absent in aerocyte ECs (aCap ECs), scale bar: 50 μm.

**C**. SPARCL1 is low / absent in large vessel ECs, including arterial (**A’’**) and venous ECs and lymphatic ECs, scale bar: 50 μm.

**Suppl Fig.3**

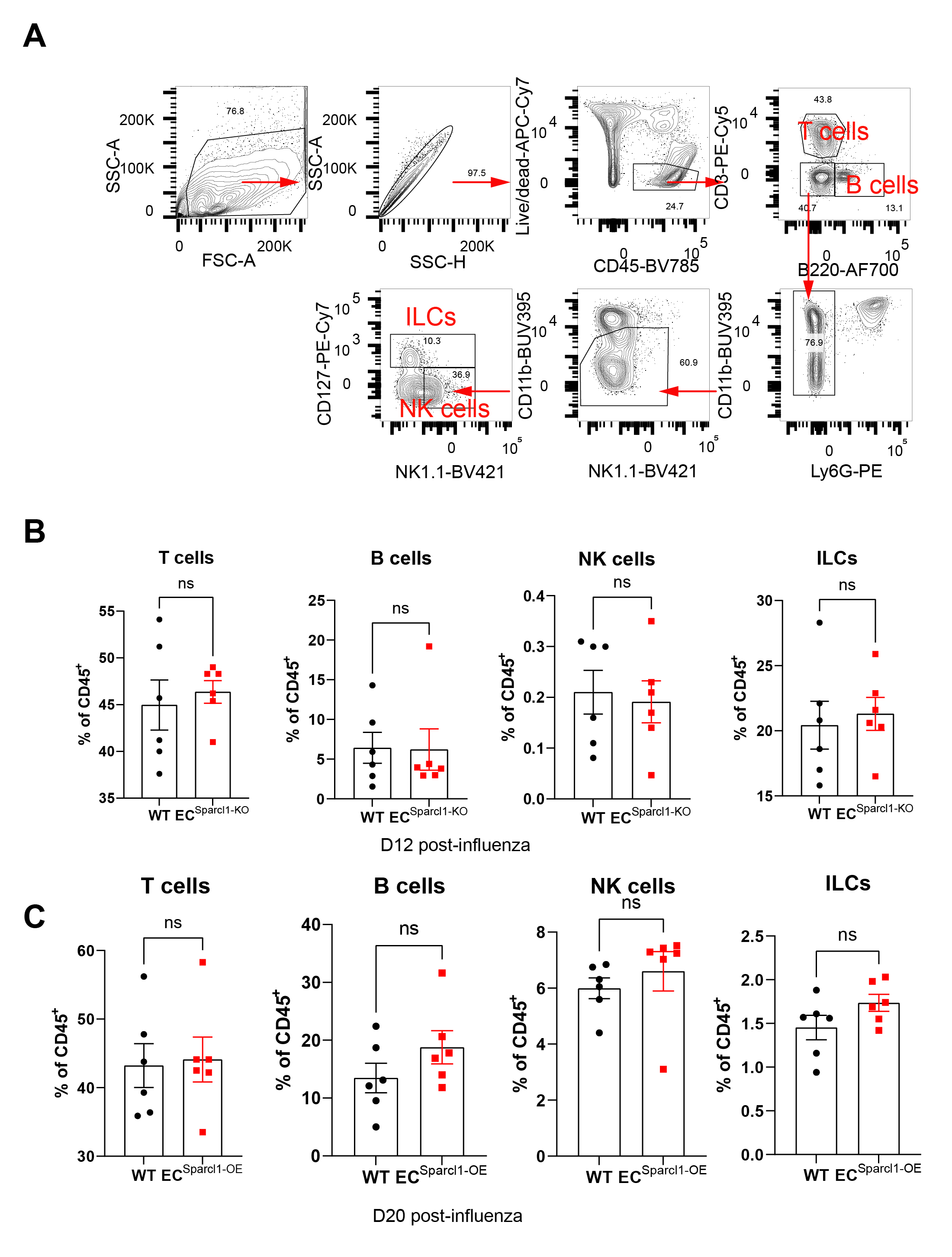

**Suppl Fig.3 SPARCL1 has no obvious effect on frequency of T, B, NK and innate lymphoid cells during influenza-induced lung injury**

**A.** Representative gating scheme for flow cytometry analysis of T, B, NK, and innate lymphoid (ILC) cells in mouse lungs after influenza injury.

**B**. Quantification of the proportion of T cells (CD3^+^B220^-^), B cells (B220^+^CD3^-^), NK cells (B220^-^CD3^-^Ly6G^-^CD11b^low/-^CD127^-^NK1.1^+^) and ILCs (B220^-^CD3^-^Ly6G^-^CD11b^low/-^CD127^+^) CD45^+^ live cells at day 12 after influenza infection in WT and EC^Sparcl1-KO^mice, n = 6 mice per group.

**C**. Quantification of the proportion of T cells (CD3^+^B220^-^), B cells (B220^+^CD3^-^), NK cells (B220^-^CD3^-^Ly6G^-^CD11b^low/-^CD127^-^NK1.1^+^) and ILCs (B220^-^CD3^-^Ly6G^-^CD11b^low/-^CD127^+^) CD45^+^ live cells at day 20 after influenza infection in WT and EC^Sparcl1-OE^mice, n = 6 mice per group.

Data in (**B**) and (**C**) are presented as means ± SEM and calculated using unpaired two-tailed t test. *P < 0.05, ns, no significance, P >0.05.

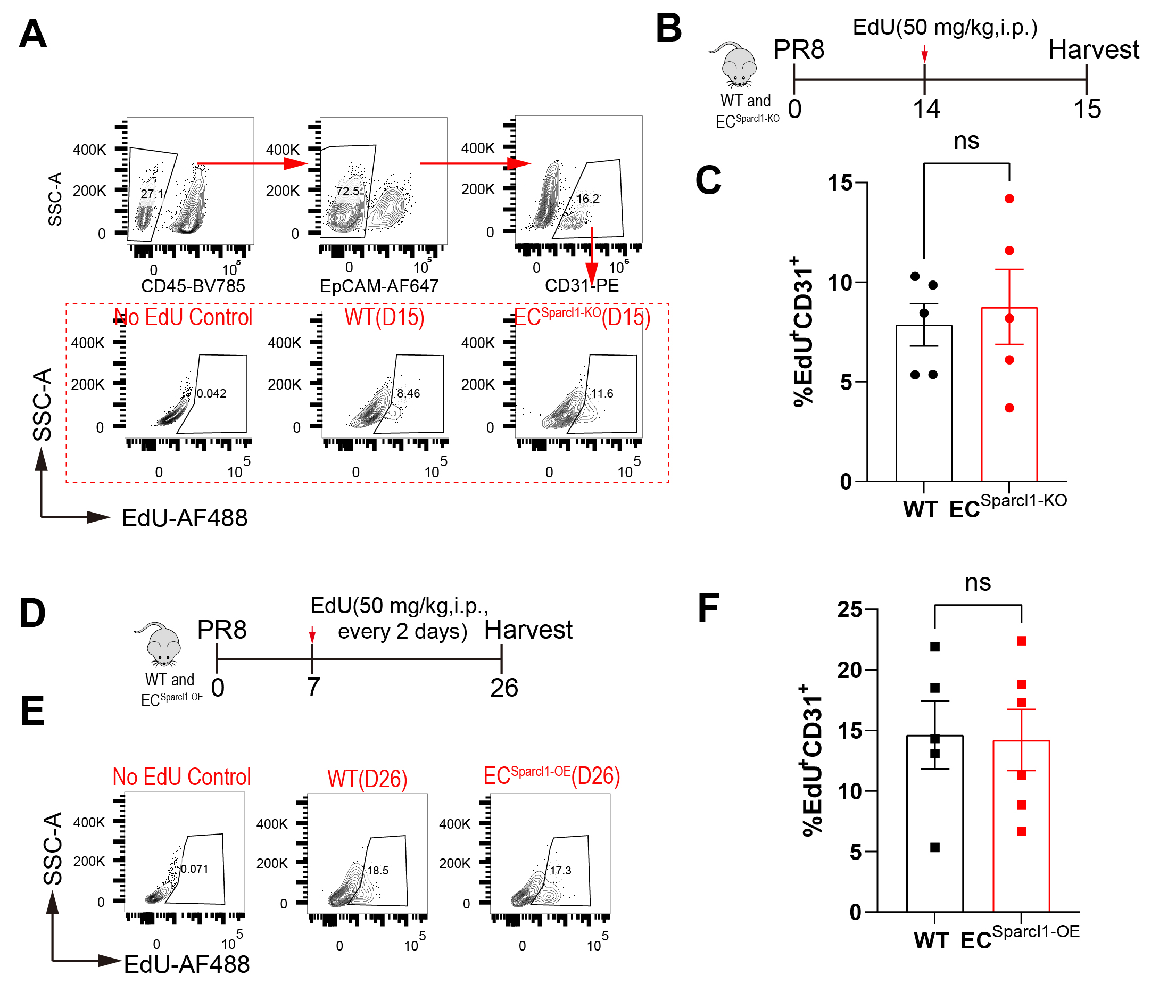
**Suppl Fig.4**

**Suppl Fig.4 SPARCL1 has no obvious effect on endothelial proliferation in influenza-induced lung injury**

**A.** Representative gating scheme for flow cytometry analysis of proliferative endothelial cells (CD45^-^EpCAM^-^CD31^+^EdU^+^) in lungs from WT and EC^Sparcl1-KO^ mice.

**B.** Timeline for EdU administration (50 mg/kg, i.p.) and sampling. WT and EC^Sparcl1-KO^ mice were treated with EdU 24 hours prior to harvest.

**C**. Endothelial EdU incorporation was measured by flow cytometry in lungs from WT and EC^Sparcl1-KO^ mice on day 15 after influenza infection, n = 5 mice per group.

**D.** Timeline for EdU administration (50 mg/kg, i.p.) and sampling. WT and EC^Sparcl1-OE^ mice were treated with EdU on day 7 after influenza infection and every 48 hours per dose later by day 26.

**E.** Representative gating scheme for flow cytometry analysis of proliferative endothelial cells (CD45^-^EpCAM^-^CD31^+^EdU^+^) in lungs from WT and EC^Sparcl1-OE^ mice.

**F**. Endothelial EdU incorporation was measured by flow cytometry in lungs from WT and EC^Sparcl1-OE^ mice on day 26 after influenza infection, n = 5 mice per group.

Data in (**C**) to (**F**) are presented as means ± SEM, calculated using unpaired two-tailed t test. *P < 0.05, **p < 0.01.

**Suppl Fig.5**

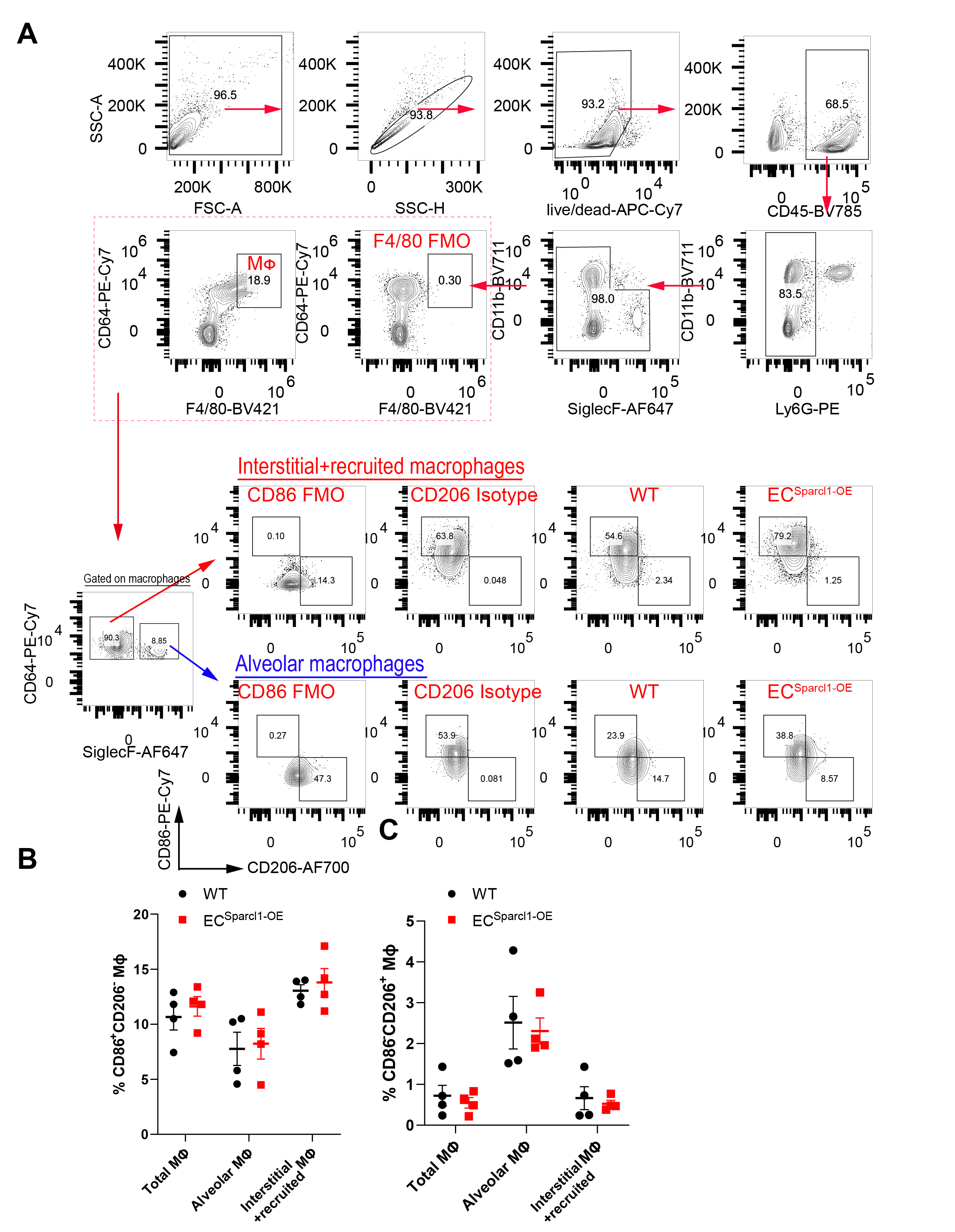

**Suppl Fig.5 Endothelial overexpression of Sparcl1 did not significantly alter the M1/M2 macrophage transition under homeostasis.**

**A.** Representative gating scheme for identification of general pulmonary macrophages (CD45^+^Ly6G^-^CD64^+^F4/80^+^) and M1-like (CD86^+^CD206^-^) and M2-like (CD86^-^CD206^+^) macrophages in total lung macrophages (CD45^+^Ly6G^-^CD64^+^F4/80^+^), alveolar macrophages (CD45^+^Ly6G^-^CD64^+^F4/80^+^SiglecF^+^) and interstitial and recruited macrophages (CD45^+^Ly6G^-^CD64^+^F4/80^+^SiglecF^-^) at day 20 after influenza infection in WT and EC^Sparcl1-OE^ mice.

**B-C.** Quantification of the proportion of (**B**)M1-like (CD86^+^CD206^-^) and (**C**)M2-like (CD86^-^CD206^+^) macrophages in total lung macrophages (CD45^+^Ly6G^-^CD64^+^F4/80^+^), alveolar macrophages (CD45^+^Ly6G^-^CD64^+^F4/80^+^SiglecF^+^) and interstitial and recruited macrophages (CD45^+^Ly6G^-^CD64^+^F4/80^+^SiglecF^-^) at day 0 after influenza infection (uninjured) in WT and EC^Sparcl1-OE^ mice, n = 4 mice per group.

Data in (**B**) to (**C**) are presented as means ± SEM, calculated using unpaired two-tailed t test. *P < 0.05.

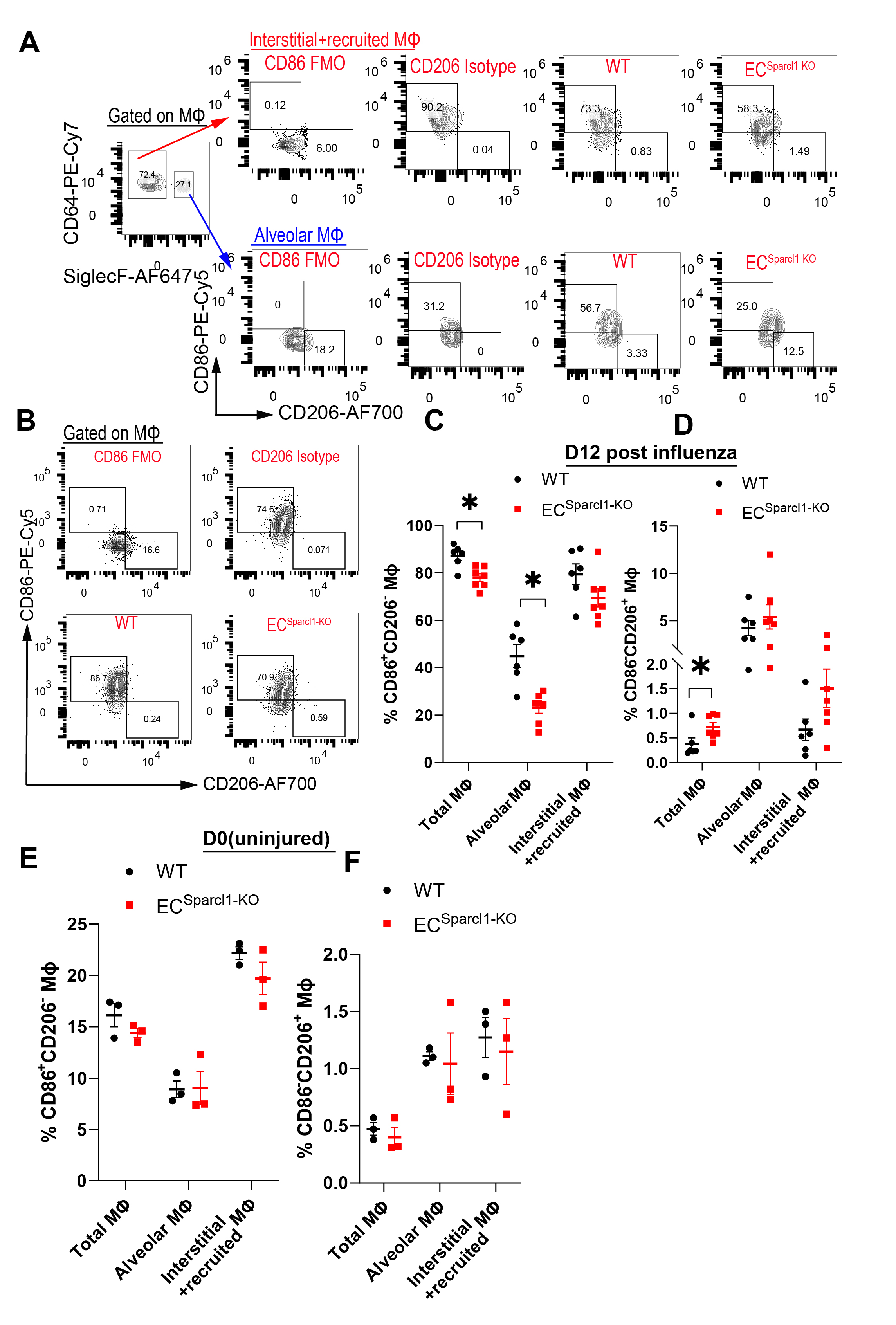
**Suppl Fig.6**

**Suppl Fig.6 Endothelial loss of Sparcl1 caused fewer M1_like but more M2_like macrophages during pneumonia**

**A.** Representative gating scheme for identification of M1-like (CD86^+^CD206^-^) and M2-like (CD86^-^CD206^+^) macrophages in alveolar macrophages (CD45^+^Ly6G^-^CD64^+^F4/80^+^SiglecF^+^) and interstitial and recruited macrophages (CD45^+^Ly6G^-^CD64^+^F4/80^+^SiglecF^-^) at day 12 after influenza infection in WT and EC^Sparcl1-KO^ mice.

**B.** Representative gating scheme for identification of M1-like (CD86^+^CD206^-^) and M2-like (CD86^-^CD206^+^) macrophages in total lung macrophages (CD45^+^Ly6G^-^CD64^+^F4/80^+^) at day 12 after influenza infection in WT and EC^Sparcl1-KO^ mice.

**C-D.** Quantification of the proportion of (**C**)M1-like (CD86^+^CD206^-^) and (**D**)M2-like (CD86^-^CD206^+^) macrophages in total lung macrophages (CD45^+^Ly6G^-^CD64^+^F4/80^+^), alveolar macrophages (CD45^+^Ly6G^-^CD64^+^F4/80^+^SiglecF^+^) and interstitial and recruited macrophages (CD45^+^Ly6G^-^CD64^+^F4/80^+^SiglecF^-^) at day 12 after influenza infection in WT and EC^Sparcl1-KO^ mice, n = 6-7 mice per group.

**E-F.** Quantification of the proportion of (**E**)M1-like (CD86^+^CD206^-^) and (**F**)M2-like (CD86^-^CD206^+^) macrophages in total lung macrophages (CD45^+^Ly6G^-^CD64^+^F4/80^+^), alveolar macrophages (CD45^+^Ly6G^-^CD64^+^F4/80^+^SiglecF^+^) and interstitial and recruited macrophages (CD45^+^Ly6G^-^CD64^+^F4/80^+^SiglecF^-^) at day 0 after influenza infection(uninjured) in WT and EC^Sparcl1-KO^ mice, n = 3 mice per group.

Data in (**C**) to (**F**) are presented as means ± SEM, calculated using unpaired two-tailed t test. *P < 0.05.

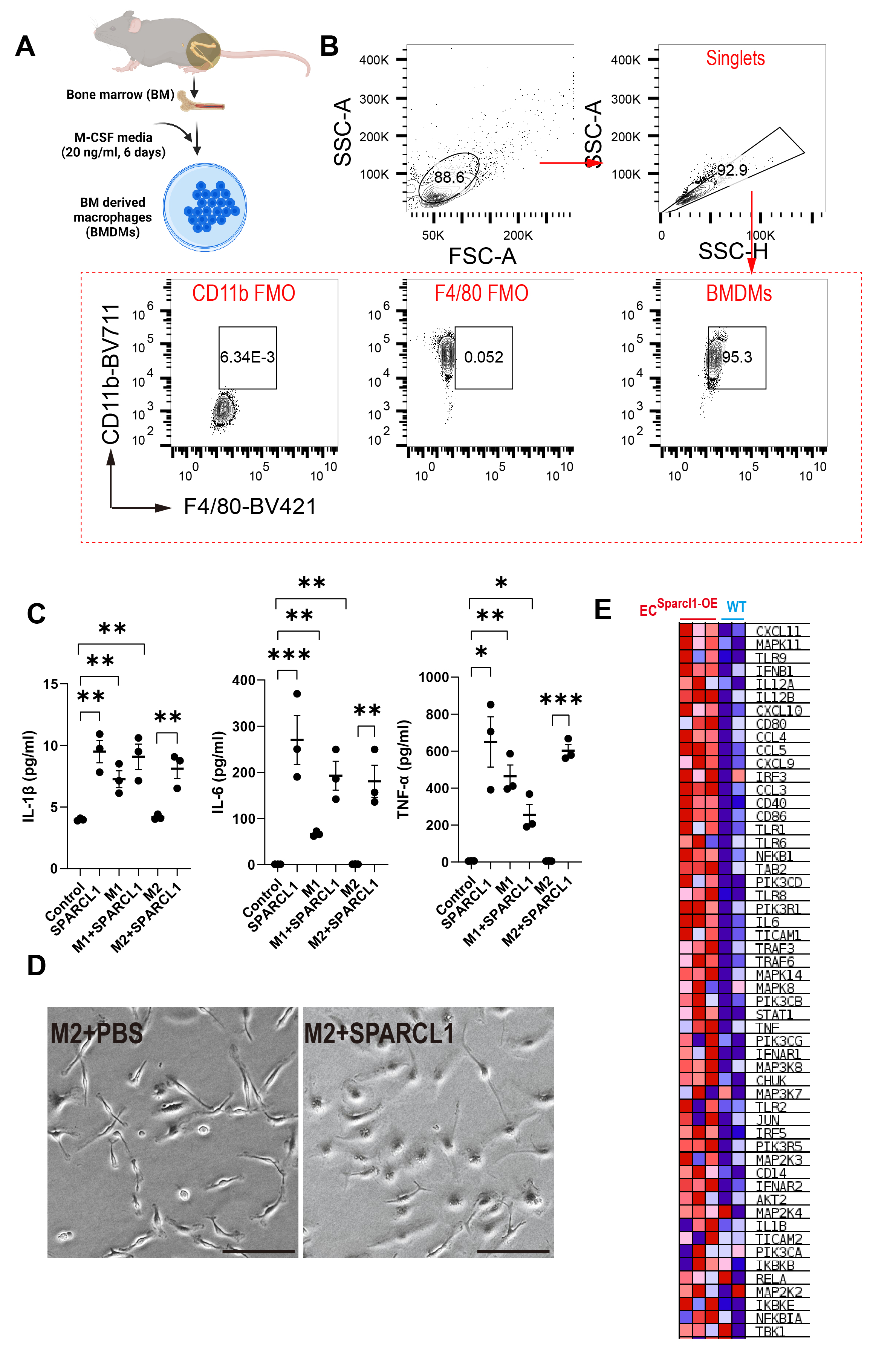
**Suppl Fig.7**

**Suppl Fig.7 SPARCL1 induces an M1-like macrophage phenotype in BMDMs**

**A.** Schematic for differentiation of bone marrow cells into bone marrow-derived macrophages (BMDMs).

**B**. FACS analysis of BMDM cell suspension. Over 95% of the cells were positive for CD11b and F4/80, which are markers for murine macrophages(*4*).

**C**. ELISA quantification of Th1 cytokines IL-1β, IL-6 and TNF-α levels in supernatant collected from M1 and M2 polarized BMDMs after treatment with SPARCL1(10 μg/ml) for 24 hours, n = 3 per group.

**D**. Cell morphology was observed 24 h after treatment of M2-polarized BMDM with SPARCL1 (10 μg/ml). Scale bar: 100 μm.

**E.** Heatmaps for genes in the TLR4 signaling pathway that positively correlated to endothelial Sparcl1 expression are shown.

Data in (**C**) are presented as means ± SEM, calculated using unpaired two-tailed t test. *P < 0.05.

**Suppl Fig.8**

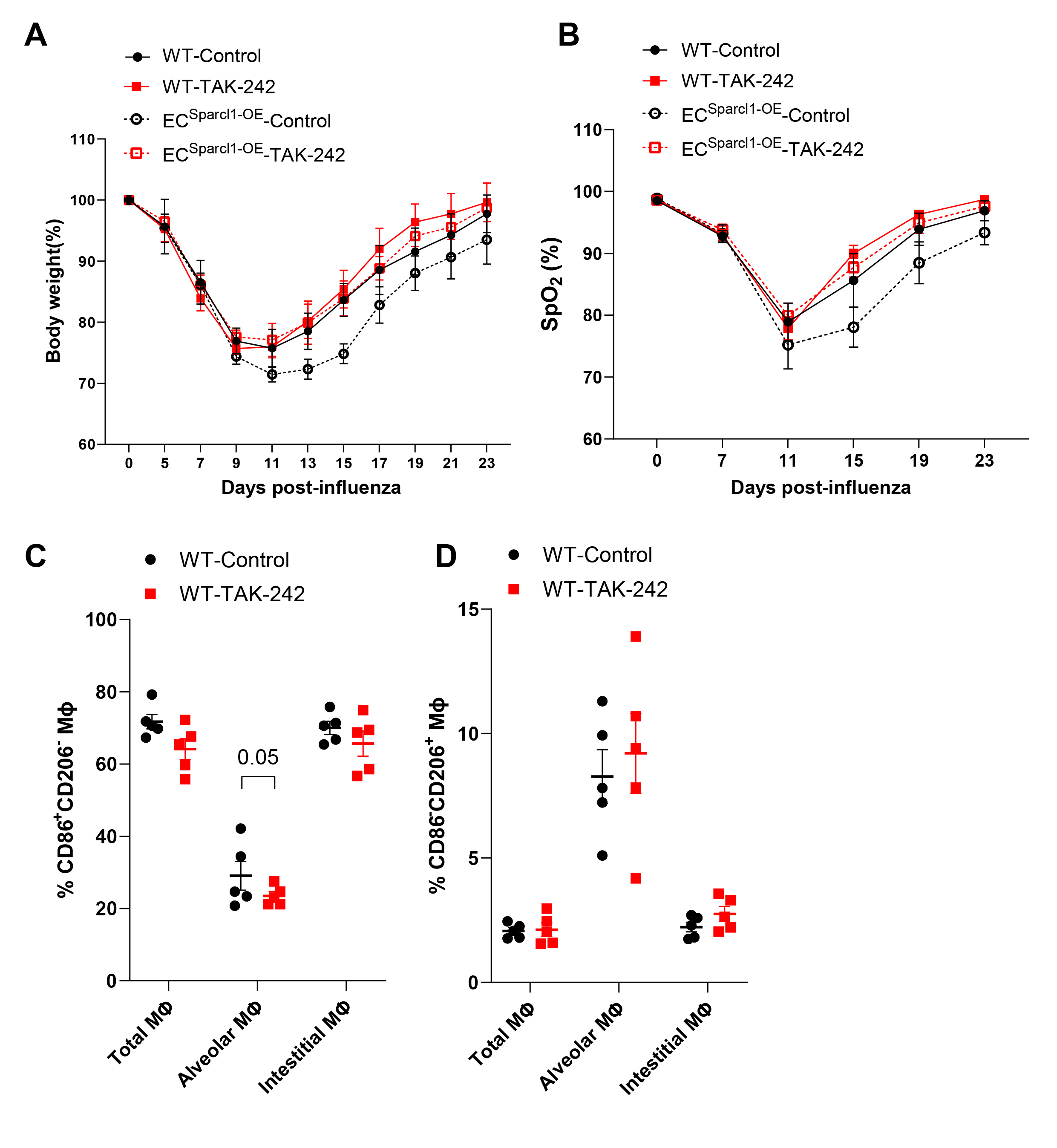

**Suppl Fig.8 TAK-242 treatment did not significantly improve influenza-induced pneumonia in WT mice**

**A-B.** Time course of changes in (**A**) body weight and (**B**) capillary oxygen saturation in WT and EC^Sparcl1-OE^ mice treated with or without TAK-242 after influenza infection, n = 5 mice per group.

**C-D**. Quantification of the proportion of (**C**)M1-like (CD86^+^CD206^-^) and (**D**)M2-like (CD86^-^CD206^+^) macrophages in total lung macrophages (CD45^+^Ly6G^-^CD64^+^F4/80^+^), alveolar macrophages (CD45^+^Ly6G^-^CD64^+^F4/80^+^SiglecF^+^) and interstitial and recruited macrophages (CD45^+^Ly6G^-^CD64^+^F4/80^+^SiglecF^-^) at day 20 after influenza infection in WT mice treated with or without TAK-242, n = 5 mice per group.

Data in (**A**) and (**D**) are presented as means ± SEM, calculated using unpaired two-tailed t test. *P < 0.05.
